# Cellular perception of growth rate and the mechanistic origin of bacterial growth law

**DOI:** 10.1101/2021.10.16.464649

**Authors:** Chenhao Wu, Rohan Balakrishnan, Nathan Braniff, Matteo Mori, Gabriel Manzanarez, Zhongge Zhang, Terence Hwa

## Abstract

Bacteria organize many activities according to their grow rate via the ppGpp signaling system. Yet it is not clear how this signaling system “knows” how fast cells grow. Through quantitative experiments, we show that ppGpp responds inversely to the rate of translational elongation in *E. coli*. Together with its roles in inhibiting ribosome biogenesis and activity, ppGpp closes a key regulatory circuit that enables the cell to perceive and control the rate of its growth across conditions. The celebrated linear growth law relating the ribosome content and growth rate emerges as a consequence of keeping a supply of ribosome reserves while maintaining elongation rate in slow growth conditions. Further analysis suggests the elongation rate itself is detected by sensing the ratio of dwelling and translocating ribosomes, a strategy employed to collapse the complex, high-dimensional dynamics of the molecular processes underlying cell growth to perceive the physiological state of the whole.

---

In the past decade, much efforts have been devoted towards characterizing and understanding the allocation of bacterial proteome across different growth conditions^1–6^. Central to the bacterial proteome allocation strategy is the approximate linear increase of the ribosome content with growth rate^7–9^, when growth is varied by using different nutrients. This classic bacterial “growth law” is rationalized by Maaloe in term of the need of more ribosomes to synthesize proteins to achieve faster growth rate, when the rate of translational elongation by ribosomes is saturated^9^. This strategy of producing ribosomes “as needed” in different growth conditions forms the basis of the optimal resource allocation strategy, which posits that cells allocate its resources (the proteome in this case) in such a way to maximize its growth^5,10^. However, it is actually long known that the translational elongation rate itself varies across growth conditions^11^, which poses a challenge to the rationalization by Maaloe. Moreover, it is known that in conditions where cells are hardly growing, a significant pool of ribosome (the “ribosome” reserves) is kept idle, presumably for rapid transition to fast growth when favorable growth conditions returns^12–14^. Intriguingly, the ribosome reserve kept by cells is not limited to slow growth, but maintained at a constant amount above the minimum needed across growth rates^2,13,15^. This behavior again challenges the notion of optimal resource allocation for the current growth condition.

One approach towards understanding the bacterial proteome allocation strategy is to follow its regulatory mechanisms, to see how the linear growth law is implemented mechanistically. This involves the sensing and control of the cell’s growth rate, since proteome allocation strategy is strongly dependent on the growth rate. Towards this end, we note that Guanosine tetraphosphate (ppGpp)^16^ is a key signaling molecule involved in bacterial response to environmental changes and in coordinating growth-rate dependent responses^17–20^. ppGpp signaling has been extensively studied^18–24^, both for mechanisms contributing to its synthesis and degradation, and for its downstream effects on hundreds of genes^25–28^, including the synthesis^29,30^ and activity^19,23^ of the translation machinery. Without a doubt, ppGpp-signaling plays a crucial role in responding to the cellular growth rate. Yet, despite the wealth of information at the molecular level, quantitative understanding of how bacteria perceive the state of cell growth is lacking. Here we reveal the underlying signaling strategies employed by *E. coli* to perceive and respond to growth, established through a series of experiments in which ppGpp and other key physiological variables are quantitatively measured. These strategies provide important insight on the initial question on bacterial proteome allocation strategy as we will discuss at the end.

## RESULTS

### A simple, robust relation between ppGpp and translational elongation rate

During environmental changes such as diauxic shifts, *E. coli* responds by producing ppGpp^31^. Fig. 1a shows a typical diauxic growth curve in minimal medium containing glycerol and a small amount of glucose as the only carbon sources: cells grow exponentially on glucose without utilizing glycerol until glucose is depleted^32^, followed by a period of growth arrest (approximately 40-50 min in this case), before fully resuming growth on glycerol. We followed the kinetics of ppGpp accumulation by performing such growth transition experiments in the presence of ^32^P-orthophosphate. Throughout the transition, labelled nucleotides were extracted and resolved by thin-layer chromatography (Fig. 1b). The ppGpp level relative to that of steady-state growth in glucose, denoted as *g*(*t*), increased by over 8-fold within the first 10 min of glucose depletion before relaxing to a new steady-state level (Fig. 1c).

**Figure 1:**
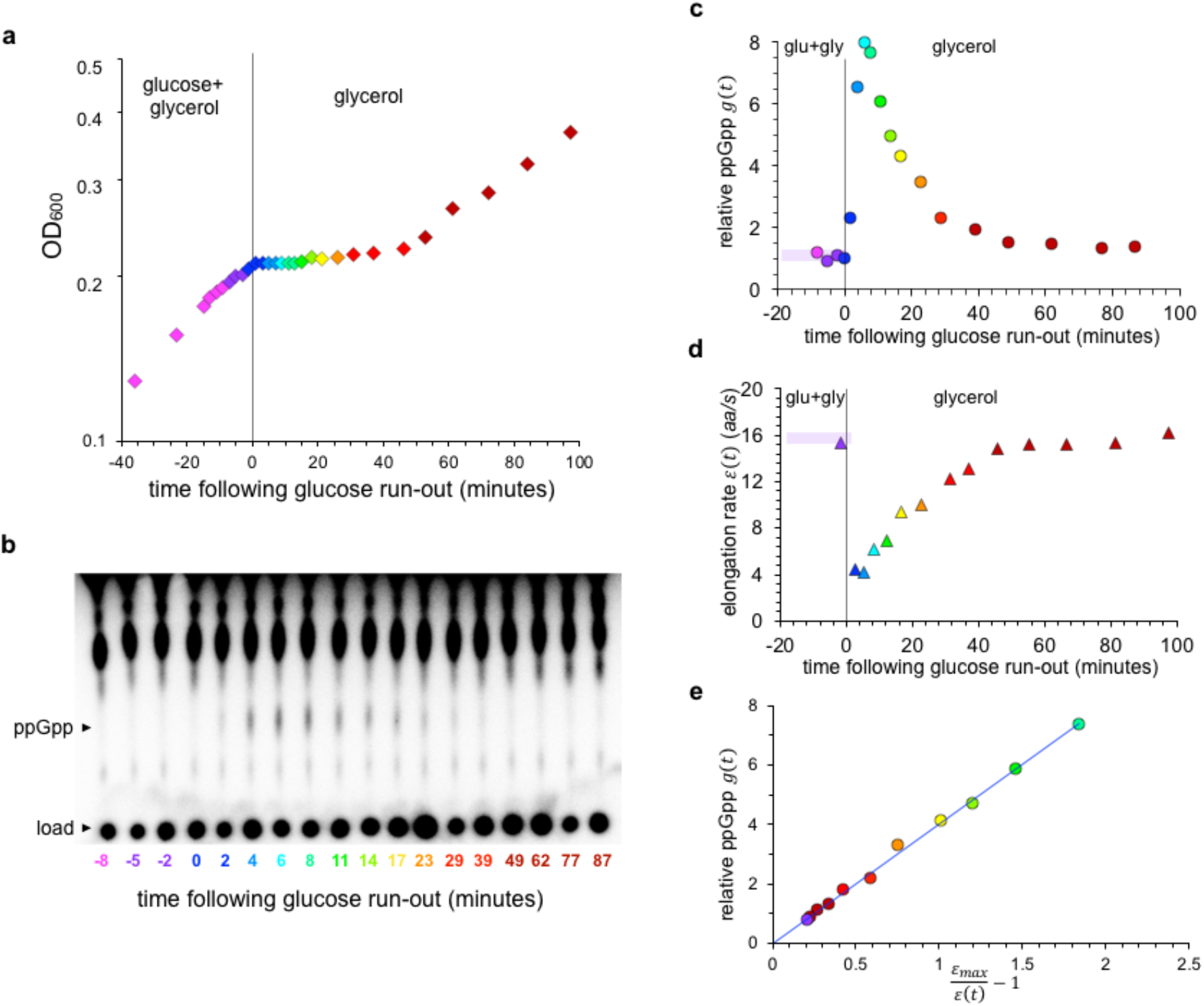
Relation between ppGpp and the translational elongation rate during growth transition. **a**, Growth kinetics of *E. coli* K-12 NCM3722 monitored by measuring the optical density at 600 nm (OD_600_) during the diauxic transition from glucose to glycerol. The same color-scheme is used across the panels to match different measured quantities to samples taken at different time during the growth transition. **b**, Resolution of ppGpp in cells sampled at different time during the growth transition by thin layer chromatography (TLC). The spots at bottome correspond to sample loading and the migrated ppGpp are indicated alongside the chromatogram. Signal intensity from the chromatograph was used as the measure of the ppGpp level. **c**, Signal intensity for ppGpp obtained from panel b is normalized by that before the growth shift, denoted by *g*(*t*), is plotted against the time *t* at which the sample was taken. **d**, The translational elongation rate *ε*(*t*) was obtained as described in Supplementary Fig. 1, and plotted against the time *t* at which sample was taken. **e**, ppGpp levels *g*(*t*) are plotted against the reciprocal of the corresponding elongation rates normalized by the maximum elongation rate (*ε*_*max*_), defined as the value of the elongation rate extrapolated to *g* = 0 (see Supplementary Fig. 1e). The line is the best-fit of the data to Eq. (1), with *c* ≈ 4.0 and *ε*_*max*_ = 19.4 ± 1.4 aa/s.

We also characterized changes in the translational elongation rate (ER, denoted by *ε*(*t*)) during the growth recovery period by assaying for the delays in LacZ induction, as previous studies have established that ER determined from LacZ is representative of that of typical proteins^33,34^, and single-molecule study of translation kinetics *in vivo* suggested little heterogeneity in ER across codons and mRNAs in the absence of antibiotics^35,36^. Using the induction time obtained at various time *t* (Supplementary Fig. 1a-c) and taking into account of the initiation time which showed little variation (Supplementary Fig. 2), the instantaneous ER, *ε*(*t*), was deduced throughout the transition period (Fig. 1d, Supplementary Fig. 1d). The data shows an abrupt drop in ER immediately following glucose depletion and a gradual recovery before growth resumed. During the period of growth arrest, the time course of *ε*(*t*) strikingly mirrored that of the relative ppGpp level *g*(*t*) (compare Fig. 1c, 1d). Scatter plot of the ppGpp level with the reciprocal of ER exhibits a striking linear relation (Supplementary Fig. 1e). Defining the value of the extrapolated ER at *g* = 0 to be *ε*_*max*_, the maximum elongation rate (to be justified below), the empirical relation between the relative ppGpp level and ER can be expressed as

**Figure 2:**
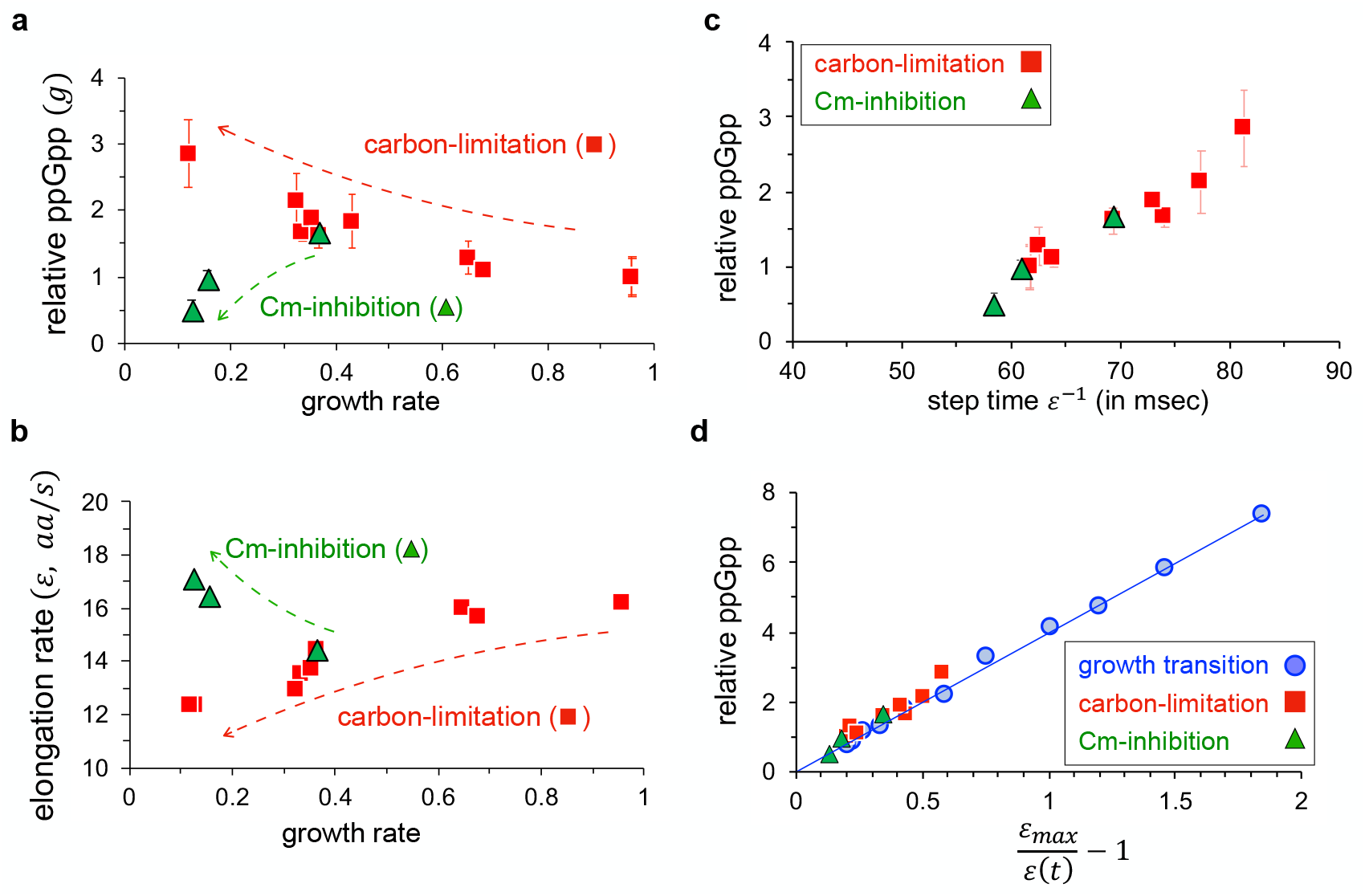
Relation between ppGpp and translational elongation rate during steady-state growth. **a**, ppGpp levels relative to that in the glucose minimal medium, *g*, are plotted against the corresponding growth rates for cells under steady state carbon-limited growth (red) and translation-limited growth (green) for cells treated with sub-lethal doses of chloramphenicol (Cm); see Supplementary Table 1. For each growth medium, ppGpp level was obtained by measuring 4 samples taken from exponentially growing cells at different ODs and using linear regression (Methods and Supplementary Fig. 13). Error bars represent the uncertainty in the linear fit. **b**, Translation elongation rates are plotted against the steady state growth rates for carbon-limited (red) and Cm treated (green) cells. **c**, Scatter plot of reciprocal elongation rates (or the step time for ribosome advancement) in milliseconds and the relative ppGpp levels (*g*) measured during steady-state growth for wild-type *E. coli* under carbon limitation (red) and translation inhibition (green). **d**, The same measurements from steady-state growth (panel c) are replotted by normalizing the elongation rate to *ε*_*max*_ together with the data collected under growth transition from Fig. 1e (blue symbols) for comparison. The line is the same as that in Fig. 1e.

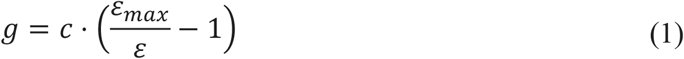

where *ε*_*max*_ ≈ 19.4 *aa*/*s* and *c* ≈ 4.0; see Fig. 1e.

Since the ppGpp level and ER are also known to change with the cellular growth rate during exponential growth^33,37^, we further examined their mutual relationship during steady exponential growth. We grew *E. coli* with different nutrient composition at growth rates ranging from 0.13 h^-1^ to 0.96 h^-1^ (Supplementary Table 1) and measured the steady-state ppGpp levels relative to that in glucose, as well as the corresponding translational elongation rates; see Methods. As growth rate was reduced, ppGpp levels increased while ER decreased, (Fig. 2a, 2b, red squares), consistent with earlier reports^33,37^ Additionally, ER has recently been shown to increase in the presence of sub-lethal amounts of chloramphenicol (Cm)^33^. Accordingly, we observed ppGpp levels to decrease and ER to increase during steady-state growth in the presence of increasing doses of Cm (Fig. 2a, 2b, green triangles). Owing to the difficulty of detecting low ppGpp levels, we used a *ΔptsG* strain (NQ1261)^33^ which has reduced glucose intake and thereby shows elevated ppGpp levels in the absence of Cm. This strain allowed us to quantify changes in ppGpp level and compare it to the changes in ER under Cm treatment. Scatter plot of the steady state ppGpp level with the reciprocal of ER under carbon limitation again exhibited a linear relation (red squares, Fig. 2c). Moreover, those from Cm-inhibited cells fell on the same linear relationship (green triangles). Strikingly, this is the same relationship as the one observed during the diauxic shift (compare with blue circles in Fig. 2d), i.e., Eq. (1) with the same intercept and proportionality constant.

### Regulatory circuit mediated by translation rate links ppGpp quantitatively to growth rate

A steady state relationship between ER and ppGpp level allows the cell to link the ppGpp level uniquely to the steady state growth rate via a simple regulatory circuit (Box 1): Due to negligible rate of protein turnover^38,39^, the rate of protein synthesis is given by the product of ER and the total number of active (translating) ribosomes per cell, 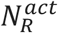. During exponential growth at rate *λ*, the total peptide synthesis rate is *λ* ⋅ *M*_*p*_, where *M*_*p*_ is the total protein mass per cell (in unit of the mass of an amino acid). The number of active ribosomes is the difference between the total number of ribosomes per cell (*N*_*R*_) and the number of inactive ribosomes per cell 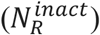. Thus,

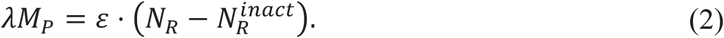

The ratio *N*_*R*_/*M*_*p*_, which is proportional to the cellular ribosome concentration (since *M*_*p*_ is proportional to the cell volume^14^), is set by the ppGpp level through regulation of rRNA expression^30^. We take this regulatory function to be

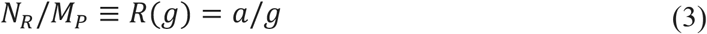

(with an unknown constant *a*) since the RNA/protein mass ratio, which is proportional to *R*, scales linearly as 1/*g*; see Fig. 3a. The inactive ribosome concentration, which is proportional to 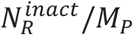, is more difficult to quantify directly due to a multitude of ppGpp-dependent effects, e.g., the binding of ribosomes to various ribosome hibernation factors including Rmf, Hpf, and RaiA^40–42^, which all increase linearly with the ppGpp level as growth rate is reduced upon limiting carbon uptake (Supplementary Fig. 3). We describe this effect by the form

**Figure 3.**
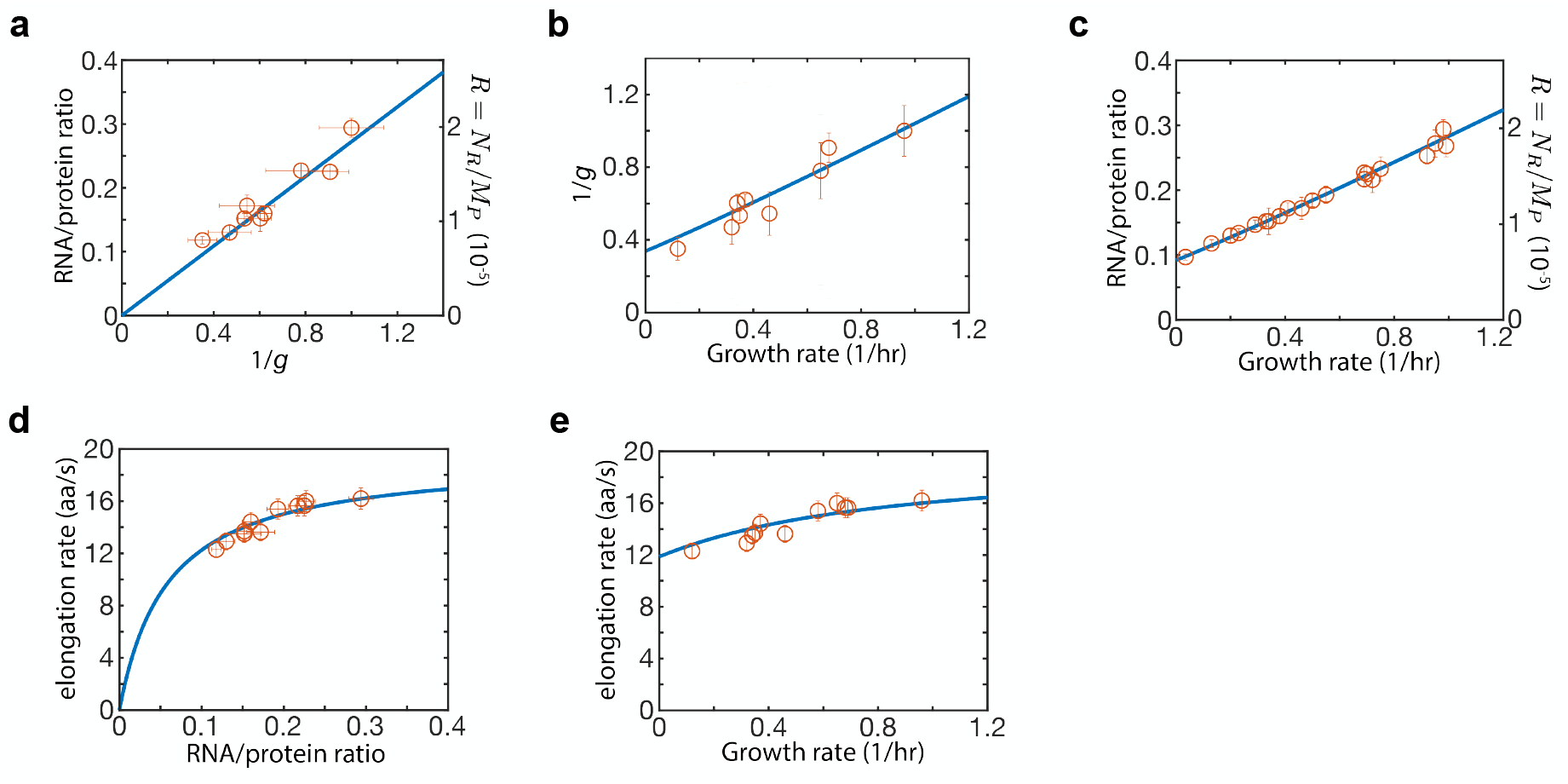
Model data comparison. **a**, RNA-protein ratio (circles, left vertical axis) is proportional to the reciprocal of ppGpp level. This ratio is taken to be proportional to the ribosome content, *N*_*R*_/*M*_*p*_ (right vertical axis), and is used as an input to the model; see Eq. (3). The proportionality constant between RNA-protein ratio and *N*_*R*_/*M*_*p*_, denoted by *η*, is one of the 3 fitting parameters of the model. The best-fit is shown by the line. **b**, The approximate linearity between the RNA-protein ratio and the growth rate (circles, left vertical axis) is well accounted for by the model (line), as is the approximate linear relation between the growth rate and the reciprocal of ppGpp level (panel **c**), and the weak relation between the ER and growth rate (panel **d**), and the Michaelis relation between ER and the RNA-protein ratio (panel **e**). The model is described by Eqs. (1) and (5). The values of the best-fit parameters were *a* ≅ 1.85 × 10^−5^, *b* ≅ 2.11 × 10^−6^, *η* ≈ 6.8 × 10^−5^.

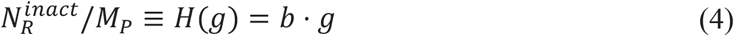

for simplicity, with another unknown constant *b*. In principle, the inhibitory effect of ppGpp on translation initiation^43–45^ would also be included in the above. However, the magnitude of this effect is not expected to be large, as no strong delay in translational initiation was detected following transient growth arrest during the diauxic shift (Supp. Fig 2).

Putting together the form of the regulatory factors in Eqs. (3) and (4) into Eq. (2) leads us to a relationship between the growth rate and the ppGpp level for exponentially growing cells:

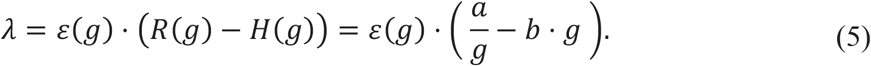

where *ε*(*g*) = *ε*_*max*_/(1 + *g*/*c*) is obtained from the steady-state relation between *ε* and *g* (Fig. 2d), which is mathematically the same as inverting Eq. (1). The two constants *a* and *b* in Eq. (5) specify the magnitudes of the two regulatory interactions. With appropriate choices of these two constants, the simple model defined by Eq. (5) is able to quantitatively capture all the observed correlations among the growth rate *λ*, the ppGpp level (*g*), the ribosome content (*R*), and the elongation rate (*ε*) under nutrient limitation (Fig. 3b-e), with model predictions based on best-fitted values of *a* and *b* shown as solid lines. In particular, the model recapitulated the well-known inverse relation between the growth rate and ppGpp level (Fig. 3b)^37,46^. This illustrates the general principle that the cell can perceive its own growth rate by incorporating the sensing of ER (Eq. (1)) into a simple regulatory circuit (Box 1) that controls the active ribosome content by the sensor. Equally importantly, the simple correspondence between the growth rate and ppGpp level enables the cell to implement growth-rate dependent control of many cellular functions, ranging from metabolism to cell division control, by simply using ppGpp to control the expression of the relevant genes^17,25,28,47,48^. In a previous study on bacterial growth control by Erickson et al^49^, an *ansatz* was introduced in which the translational activity (*σ* ≡ *λ*/*R*) was used to successfully predict gene expression dynamics during diauxic shifts. The results here establish a one-to-one relation between *σ* and *g* (Supplementary Fig. 4), thereby justifying the *ansatz* used in Ref. ^49^.

### Linear bacterial growth law obtained with a special condition on elongation rate

At a quantitative level, our model captured the approximate linear relation between the ribosome content and the growth rate (Fig. 3c), the celebrated growth law discovered long ago^7–9^. Additionally, the model captured the Michaelis-like relation between the ER and the ribosome content (Fig. 3d), substantiated with extensive data collected from many conditions as reported in Dai et al^33^. Notably, a fit of the data to the Michaelis-Menten relation (Supplementary Fig. 5) recovers a maximum elongation rate (20.0 ± 1.9 aa/s) that is indistinguishable from *ε*_*max*_ = 19.4 ± 1.4 aa/s defined by taking *g* → 0 in Eq. (1) (Supplementary Fig. 1e). Finally, the model captures the weak dependence of the elongation rate at different growth rate (Fig. 3e), with the minimal ER in slow growth condition, denoted by *ε*_0_, whose value is close to one-half of *ε*_*max*_.

The emergence of a simple linear relation between the ribosome content *R* and the growth rate *λ* is surprising, given the nonlinear regulatory effects exerted by ppGpp (Eq. (5)). In fact, a variety of relationships among these quantities is possible for generic values of *a* and *b* (Supplementary Fig. 6). However, if the regulatory parameters *a* and *b* are such that the ratio *ε*_0_ : *ε*_*max*_, is exactly one-half, then mathematically the model yields an exact linear relation between *R* and *λ*, with the slope given by 1/*ε*_*max*_, and an exact Michaelis-Menten relation between the ER and the ribosome content, with the maximal ER being *ε*_*max*_ (Supplementary Note 1). Thus, to the extent that Eqs. (3) and (4) capture the forms of the regulatory functions, prescribing the appropriate regulatory parameters *a* and *b* to enforce *ε*_0_ being approximately one-half of *ε*_*max*_ is required for the emergence of the approximate linear growth relation between the ribosome content and the growth rate. (We have separately shown that adding offsets to the simplest forms of the regulatory functions used in Eqs. (3) and (4) do not affect the quality of the fit; see Supplementary Fig. 7.)

Maintenance of ER above a minimal level is clearly of physiological importance, as too low an ER would lead to problems in the processivity of protein synthesis^50,51^. Another physiological requirement is the maintenance of a sufficient ribosome reserve at slow growth, denoted by *R*_0_, needed for rapid growth recovery when favorable nutrient conditions return^14^. Both physiological requirements are satisfied by employing hibernation factors to inactivate ribosomes. By employing both positive and negative regulation through distinct promoters (Box 1), the cell can readily attain the required values of *ε*_0_ and *R*_0_ by simply prescribing the regulatory parameters *a* and *b*; see Supplementary Fig. 8 and Supplementary Note 1. To keep *ε*_0_ high while also maintaining a ribosome reserve is possible in principle; see Supplementary Fig. 9. With high ER (e.g., above 90% of *ε*_*max*_, dashed line in Supplementary Fig. 9a), the ribosome content would even be moderately reduced at fast growth rate (Supplementary Fig. 9b), a fitness benefit from the proteome allocation perspective^2,52^. However, this strategy would also require exquisite mechanism for detecting very small changes in ER (Supplementary Fig. 9d). Thus, the choice of using *ε*_0_ ≈ *ε*_*max*_/2 may reflect a compromise between the physiological demand for keeping *ε*_0_ high and the molecular constraint for detecting small changes in ER in order to sense slow growth and enforce growth-rate dependent regulation.

### Mechanism of sensing the translational elongation rate

It is important to point out that the steady-state results presented here (Eq. (5) and Fig. 3) is predicated on the existence of the empirical relation given by Eq. (1). We now return to discuss the causal link and the mechanistic origin of this empirical relation. Towards this end, the first question to address is whether ppGpp or ER is the primary driver of this response. One scenario is that ppGpp rises in response to some unknown “starvation signal” as glucose ran out, and the resulting increase in ppGpp then reduces the ER. However, we will show shortly below that a mutant in which ppGpp does not rise instantaneously still exhibits a strong immediate drop in ER as glucose ran out. An alternative scenario is that the drop of ER occurs first, and this drop is itself the signal that drives up ppGpp. The latter scenario is supported by metabolomic study which found the amino acid pools (and particularly the glutamate pool) to drop sharply and immediately following glucose runout^53^, thus imposing an obligatory reduction in ER. Sensing the drop in ER could therefore be an effective strategy to sense the nutritional status of the cell.

We next examine the form of the response (1) in terms of the known mechanisms of ppGpp synthesis and degradation. It will be convenient to re-express ER and *ε*_*max*_ in Eq. (1) in terms of the elemental steps of the translation cycle (Box 2a): a time *τ*_*dwell*_ where the ribosome dwells on the A-site waiting for the cognate charged tRNA, and a time *τ*_*trans*_ for peptidyl transfer and translocation to the next codon. This changes Eq. (1) to *g* = *c* ⋅ *τ*_*dwell*_/*τ*_*trans*_, with the maximal elongation rate 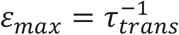 identified with the case where *τ*_*dwell*_ → 0. (Incidentally, the special limit of elongation rate at slow growth being one-half of *ε*_*max*_ corresponds to *τ*_*dwell*_ = *τ*_*trans*_ at slow growth.)

Next, detailed analysis based on flux balance (Supplementary Note 2) establishes a simple relation between two pools of actively translating ribosomes, those in the dwelling state (of concentration *R*_*dwell*_) and those in the process translocation (of concentration *R*_*trans*_):

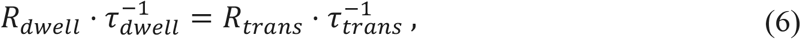

Eq. (6) is simply a condition of *detailed balance* between the flux of ribosomes transitioning from the dwelling state to the translocation state, and the flux transition from the translocation state back to the dwelling state, with *R*_*dwell*_ + *R*_*trans*_ = *R*_*act*_ being the total concentration of actively translating ribosomes. In terms of these ribosome pools, Eq. (1) then becomes

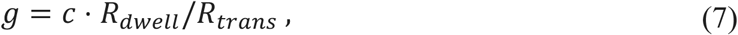

i.e., the ratio of the two pools of ribosomes.

In a simple model of ppGpp comprised of rapid equilibration between synthesis and degradation such that 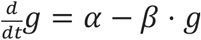, the ppGpp level is given by the ratio of the synthesis rate (*α*) and the specific degradation rates (*β*). A simple scenario giving rise to the empirical relation (1) or (7) is therefore to have the synthesis rate *α* ∝ *R*_*dwell*_ and the degradation rate *β* ∝ *R*_*trans*_. The effect of dwelling ribosomes on ppGpp synthesis is well-supported molecularly based on the known structure of the ppGpp synthetase RelA in complex with the ribosome^54,55^ as well as earlier biochemical studies^56,57^, which find that RelA is activated (i.e., synthesizing ppGpp) only when it is complexed with an uncharged tRNA together at the A-site, i.e., when the ribosome is in the dwelling state (Box 2b left). Less is known about ppGpp degradation, which is solely catalyzed by SpoT in *E. coli*^20,58^. The empirical relation (1) excludes a model with constitutive hydrolysis of ppGpp by SpoT (Supplementary Note 2 and Supplementary Fig. 10) and instead predicts regulation of SpoT hydrolysis activity, e.g., its stimulation by the translocating ribosomes (Box 2b right) or more complex regulation of SpoT activity depending on the ppGpp level itself as discussed in Supplementary Note 2. Consistent with these ideas, we find deletion of *relA* to disrupt the linear relationship between ppGpp and ER during the diauxic transition (Supplementary Fig. 11), with the remaining nontrivial ppGpp dynamics attributed to the response of SpoT, which is also the only other enzyme capable of ppGpp synthesis in *E. coli*^20,24^. [In this strain, a much slower accumulation of ppGpp occurred following glucose runout compared to the wild type (Supplementary Fig. 11b); yet ER dropped more and for a longer period, thus negating the afore-mentioned scenario that the drop of ER resulted from ppGpp accumulation.]

Intriguingly, *relA* deletion has no effect on either the ppGpp level or the elongation rate in steady state growth (Supplementary Fig. 12), consistent with the knowledge that RelA is not essential during steady-state growth^37^. The maintenance of the relationship (1) by the *ΔrelA* strain can only be attributed to SpoT. Given Eq. (7), our data thus suggest that in steady state, SpoT has acquired the ability to sense both the dwelling and translocating ribosomes, in ways that it is not capable of during transient shift. How SpoT may sense different states of the ribosome is however not known. While there is some evidence of SpoT associated with the ribosome^59^, the functional significance of this association is unclear. It is likely that this process is aided by some unknown mediators which interact with the ribosomes in ways analogous to RelA (Box 2b) and convey that information to SpoT.

## DISCUSSION

Coordination of bacterial behaviors with growth rate is widely observed^3,48,60,61^. While the ppGpp signaling system is known to be central to the growth-rate dependent responses^17,18^, how growth rate is perceived and used for regulation is not known at the quantitative level. Our work establishes a missing central element in *E. coli*’s strategy to perceive its state of growth and respond to it: Changes in nutrient environment is immediately reflected in the rate of translational elongation, and this in turn sets the ppGpp level within a time scale of 5-10 mins (Fig. 1). ppGpp’s well-established regulation of the ribosome content and activity, together with its robust relation with the elongation rate (Box 1), then forges a unique relation between ppGpp and growth rate since the latter is given quantitatively by the product of active ribosomes and their elongation rate.

As translational elongation is affected by many metabolic processes in the cell, monitoring the elongation rate is an effective strategy to diagnose the state of cell growth independently of specific nutrient bottlenecks. This is the origin of the generality of the phenomenological growth laws, i.e., why the same quantitative relation between the ribosome content and the growth rate is sustained regardless of whether cell growth is limited by carbon supply^2,7,62^, nitrogen supply^63^, partial auxotrophy^3,62^, or drugs which inhibits tRNA charging^64^. This mechanism also predicts generally that for perturbations not captured by a repartition of the two ribosomal states, including antibiotics^33,65^, phosphate limitation^63^ and lipid stress^66^, growth rate perception is distorted and the linear relation between ribosome content and growth rate is altered.

One surprising finding of our analysis is the important role played by factors that inactivate the ribosome. These factors control the amount of ribosome reserve kept by cells at slow growth^67^, and the ribosome reserve is important because it affects the rapidity of growth recovery when good growth condition returns^12–14,67^. However, the *necessity* of employing ribosome inactivating factors can only be appreciated in light of cell physiology at slow growth: If inactivating factors are not used, then keeping a finite ribosome reserve would require the translational elongation rate to drop to low levels in slow growth conditions, leading to problems in protein synthesis^68^. Thus, these inactivating factors serve as an effective tool to maintain elongation rate while setting aside ribosome reserve at slow growth.

Nevertheless, the deployment of these factors complicates the overall regulation of active ribosomes, making it difficult to understand how the well-known linear relation between ribosome content and growth rate arises. Our analysis shows that the linear relation emerges for a special choice of regulatory parameters such that the elongation rate at very slow growth is one-half of the maximal elongation rate, or alternatively, the time a ribosome spends in the dwelling state of the translation cycle is not longer than the time it spends in the translocating state, i.e., *τ*_*dwell*_ ≤ *τ*_*transl*_. It should be noted that this implementation of the linear ribosome-growth rate relation is very different from the existing ideas, based e.g., on a Michaelis-relation between translational factors and the elongation rate^15^, or on optimizing the steady-state growth rate^5,10^. Regarding the former, this work shows that the appearance of a Michaelis relation between the translational factors and the elongation rate^33^ is actually a consequence of regulation involving the ribosome inactivating factors. Regarding the latter, the existence of ribosome reserve which is detrimental to steady state growth but needed for rapid transition from slow to fast growth, is typically glossed over in optimization theories. However, the specific mechanism used to set the reserve, i.e., the use of ribosome inactivating factors which are needed for maintaining a finite elongation rate in the presence of inactive ribosomes, and which produces a constant “offset” (*R*_0_) in the linear ribosome-growth rate relation across the entire growth rate range, calls into question the popular notion that the linear ribosome-growth rate relation is predominantly driven by the optimization of steady-state growth. Instead, it suggests that setting aside a pool of ribosomes as a reserve is something specific that the cell intends to accomplish in its proteome allocation strategy, despite having a cost to the steady-state growth rate.

Turning to the mechanisms for sensing the translational elongation rate, our data and analysis show that it is based on the ratio of translating ribosomes in their two alternating states (Box 2). Sensing of the dwelling ribosomes fits well with the elaborate molecular design known for RelA^24,57^. However, our analysis suggests that this RelA-based mechanism is insufficient by itself: As RelA is not required in steady state, ppGpp synthesis activity by SpoT must somehow also be able to respond to dwelling ribosomes. The synthesis activity of SpoT was recently shown to be correlated with the levels of acetyl phosphate, a glycolytic intermediate^69^. But glycolytic flux is not necessarily a proxy for dwelling ribosomes. While SpoT has been shown to interact with ribosome associated proteins^70^, currently it remains unclear what interactions can enable SpoT to sense dwelling ribosomes. Moreover, even in the presence of RelA, the empirical relation observed between ppGpp and elongation rate during growth transition requires additional regulation by SpoT (Supplementary Note 2). Finer details of the ppGpp signaling system may be revealed by quantifying how the ppGpp-ER relation is modified for various combinations RelA and SpoT mutants in future studies.

At a broader level, this study provides a rare, trackable example of how cells perform dimensional reduction at the molecular level to attain crucial physiological information at the cellular level^71^. The key “trick” *E. coli* uses to collapse the high-dimensional complexity of the metabolic state of a cell, e.g., involving 20 amino acid synthesis pathways and the charging of over 60 tRNAs (Box 2c), is to take advantage of detailed balance between the two alternating states of the elongation ribosome, so that the ratio of the ribosome dwelling and translocation time, which reflects a weighted average of the tRNA charging ratios, can be deduced from the ratio of the dwelling and translocation pools of ribosomes regardless of molecular details (Box 2a, 2b and Supplementary Note 2). Identifying and elucidating further instances of such strategies of dimensional reduction employed by cells will be instrumental in fundamentally advancing our understanding of the connection between molecular interaction and cellular physiology for prokaryotes as well as for eukaryotes.

## Methods and Materials

### Growth media composition and culture conditions

Steady-state and growth transitioning cultures were grown in MOPS based minimal media^72^ supplemented with various carbon sources and chloramphenicol as indicated in Supplementary Table 1. All cultures were grown at 37°C with shaking at 250 rpm. For every experiment, culturing was carried out through sequential propagation of seed cultures in LB, pre-cultures in the experimental medium, and the experimental cultures. Single colonies from fresh LB agar plates were first grown in LB broth for 6 hrs as the “seeding culture”. In the pre-culturing step, depending on the experiment, cells from seeding cultures were diluted into appropriate media such that the pre-cultures would still be in exponential growth phase after overnight growth. Media used for pre-culturing in steady-state experiments were same as the experimental media (Supplementary Table 1). For the glucose to glycerol transitions, optical density was monitored at 600 nm (OD_600_) to follow the growth transition kinetics. Pre-cultures were grown in medium supplemented with 20mM glucose and 20mM glycerol, to avoid glucose run-out during the pre-culturing step. Exponentially growing pre-cultures were then diluted in the appropriate experimental medium (pre-warmed) at an initial OD_600_ of ∼0.005 and various measurements were carried in the OD_600_ range of 0.1-0.4.

### Strain construction

Wild type *E. coli* K-12 NCM3722^73,74^ and its derivatives were used in this work. HE838 *(ΔrelA*) was constructed using the λ-Red recombinase method^75^ as follows. The km resistance gene was amplified from pKD13 using chimeric oligos relA1-P1 and relA2-P2 (Supplementary Table 2). The PCR products were electroporated into NCM3722 cells expressing Lambada-Red proteins encoded by pKD46. The Km resistant colonies were confirmed by PCR and sequencing for the replacement of the region harboring *relA* by the Km gene.

### Translation elongation rate (ER) measurement

ER was measured using LacZ as a reporter as described in Dai et al^33^ with modifications. Depending on the experiment, 10 ml cultures were either grown in different steady state conditions or as undergoing glucose to glycerol growth-transition. When cultures reached OD_600_ = 0.4 (for steady-state growth) or at specific time-points during the growth transition, 5 mM isopropyl-β-D-thiogalactoside (IPTG) was added to induce the *lac* operon. Immediately after induction, 500 ml samples were taken at 10 s or 15 s intervals to pre-cooled (−20°C) tubes with 20 ml of 0.1M chloramphenicol and then rapidly frozen by dry ice. Samples were stored at -80°C before beta-galactosidase assay. 4-methylumbelliferyl-D-galactopyranoside (MUG, a sensitive fluorescence substrate) was used to measure LacZ activities in this work. Briefly, each sample was diluted by Z-buffer by 5-fold and added to 96-well plate to a volume of 200 ml. Plate was warmed at 37°C for 10 mins before adding MUG. Tecan (SPARK) plate reader was used for MUG injection and fluorescence readings. 20 ml of 2mg/ml MUG was injected to each well and fluorescence intensity (365nm excitation filter, 450nm emission filter) was measured every 4 mins for 2 hours. In the linear range of fluorescence intensity vs time plot, a linear fit was applied to obtain the slope as the relative LacZ activity for each sample. By plotting the square root of the relative LacZ activity above basal level against time^76^ (Schleif plot), the lag time for the synthesis of the first LacZ molecule (*T*_*first*_) was obtained for each sample; see Supplementary Fig. 1a-c. Similar measurements using the *α*-complement of LacZ^33^ (strain NQ1468) allows us to estimate the translational initiation time; see Supplementary Fig. 2. The ER measured in this work was found to be slightly higher than that reported by Dai et al^33^, likely due to the higher sensitivity of the substrate MUG compared to ONPG as used previously^33^.

### ppGpp measurement

ppGpp measurements were carried out as described by Cashel^77^ with minor modifications. Typically, experimental cultures were grown in 3ml volumes. Labelling was carried out when the experimental cultures grew to OD_600_ = 0.02 by adding 0.1mCi ^32^P-orthophosphate (Perkin Elmer) per ml culture. For steady-state growth, 20 μl aliquots were drawn at various OD_600_ values between the range 0.1-0.4 (see Supplementary Fig. 13a), and added to an equal volume of pre-chilled 10% formic acid. For cultures undergoing diauxic shift, 20 μl aliquots were drawn at various time points during the transition, and added to an equal volume of pre-chilled 10% formic acid. Formic acid-extracts were spun down at 13k rpm for 10 minutes and a total of 2 μl supernatant was spotted 0.5 μl at a time near the base of a PEI-Cellulose F thin layer chromatography plate (Millipore). The spots were dried and nucleotides were resolved using freshly prepared 1.5M KH_2_PO_4_ (pH 3.4). The TLC plates were dried and exposed to a phosphorimaging screen for 24-36 hours. Chromatograms were imaged using a Typhoon FLA 9500 scanner (GE) and analyzed using Fiji software. For steady state growth conditions, the slope of ppGpp signal intensities versus OD600 were compared among different cultures to obtain the relative ppGpp levels, as shown in Supplementary Fig. 13c. For any batch of measurements, the ppGpp level from a sample of NCM3722 growing steadily in MOPS glucose was always included as a reference. All measurements in that batch were normalized to the glucose-grown reference in the same batch.

### Total RNA and Protein measurement

Total RNA was measured using the method of Benthin et al^78^, and protein was measured using the Biuret method^79^, with minor modifications as described by You et al^62^.

## Acknowledgements

We are grateful to technical advices from Xiongfeng Dai and Manlu Zhu in early stages of this work. We also thank helpful discussions with Mike Cashel on the general roles of SpoT, with Frank Bruggeman on models of ppGpp dynamics and with Boris Shraiman on the general issue of dimension reduction in biological regulation. This work was supported by the NSF through Grant MCB-1818384 and the NIH through Grant GM109069.

---

**Box 1:**
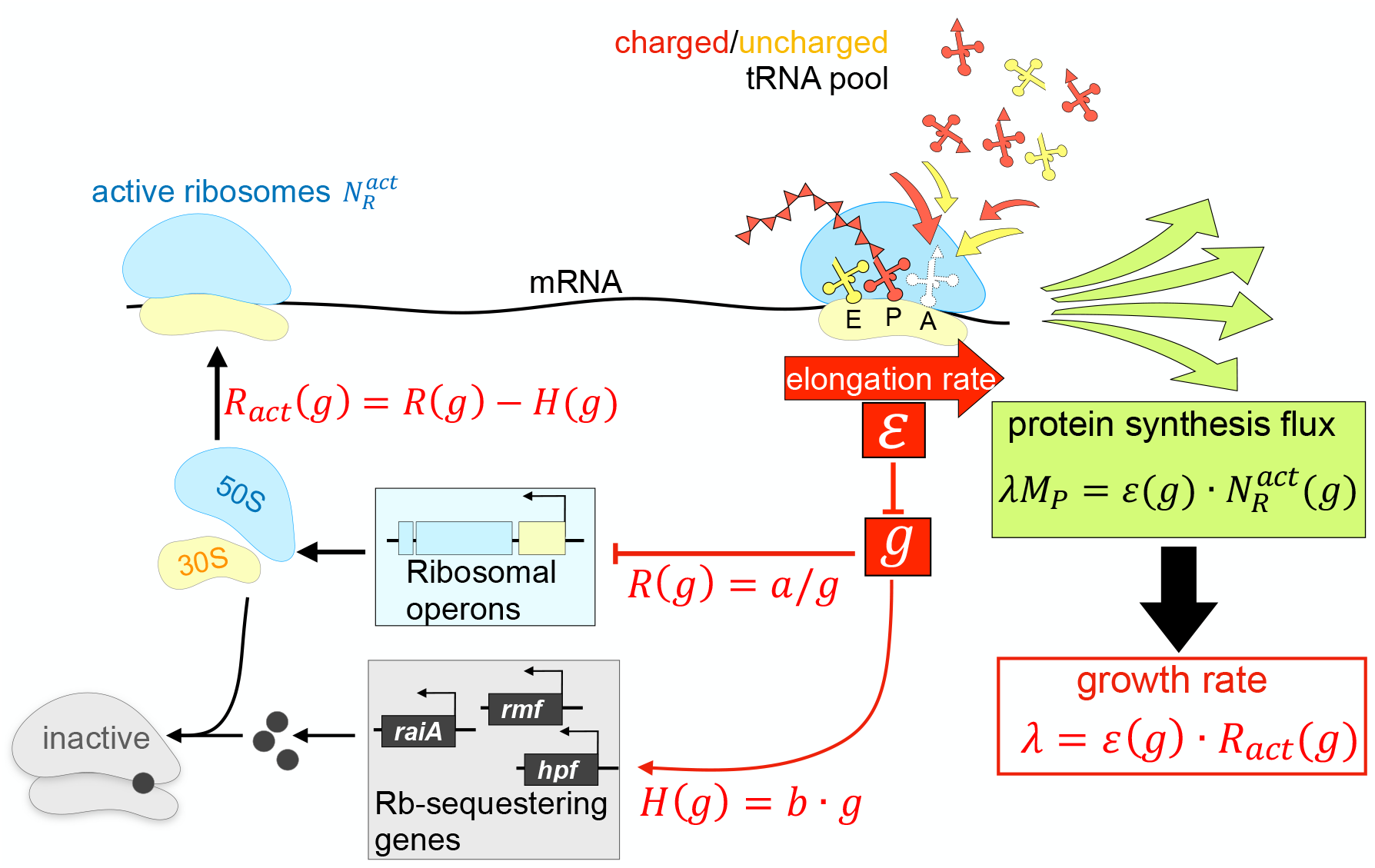
Regulatory circuit connecting ppGpp to growth rate. The steady state protein synthesis flux *λ* ⋅ *M*_*p*_ is given by the product of the elongation rate *ε* and the number of active ribosomes, 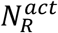. Because *ε* is simply connected to the ppGpp level *g* (Fig. 2d as summarized by Eq. (1)), and the active ribosome content 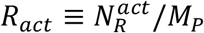 is given by the difference between the total ribosome content *R*(*g*) (Fig. 3a and Eq. (3)) and the content of the ribosome-sequestering elements *H*(*g*) (Supplementary Fig. 3 and Eq. (4)), each of which is a function of *g* due to ppGpp-mediated regulation, it follows that *λ* = *ε*(*g*) ⋅ *R*_*act*_(*g*) is a function of *g*. This gives rise to the correlation between the ppGpp level and growth rate^37,46^(Fig. 3b). By using ppGpp to regulate a spectrum of cellular processes^19,22,24^, the cell thus manages to link the regulation of these processes to the growth rate, leading to the appearance of “growth-rate dependent” control^17,47,60^.

**Box 2:**
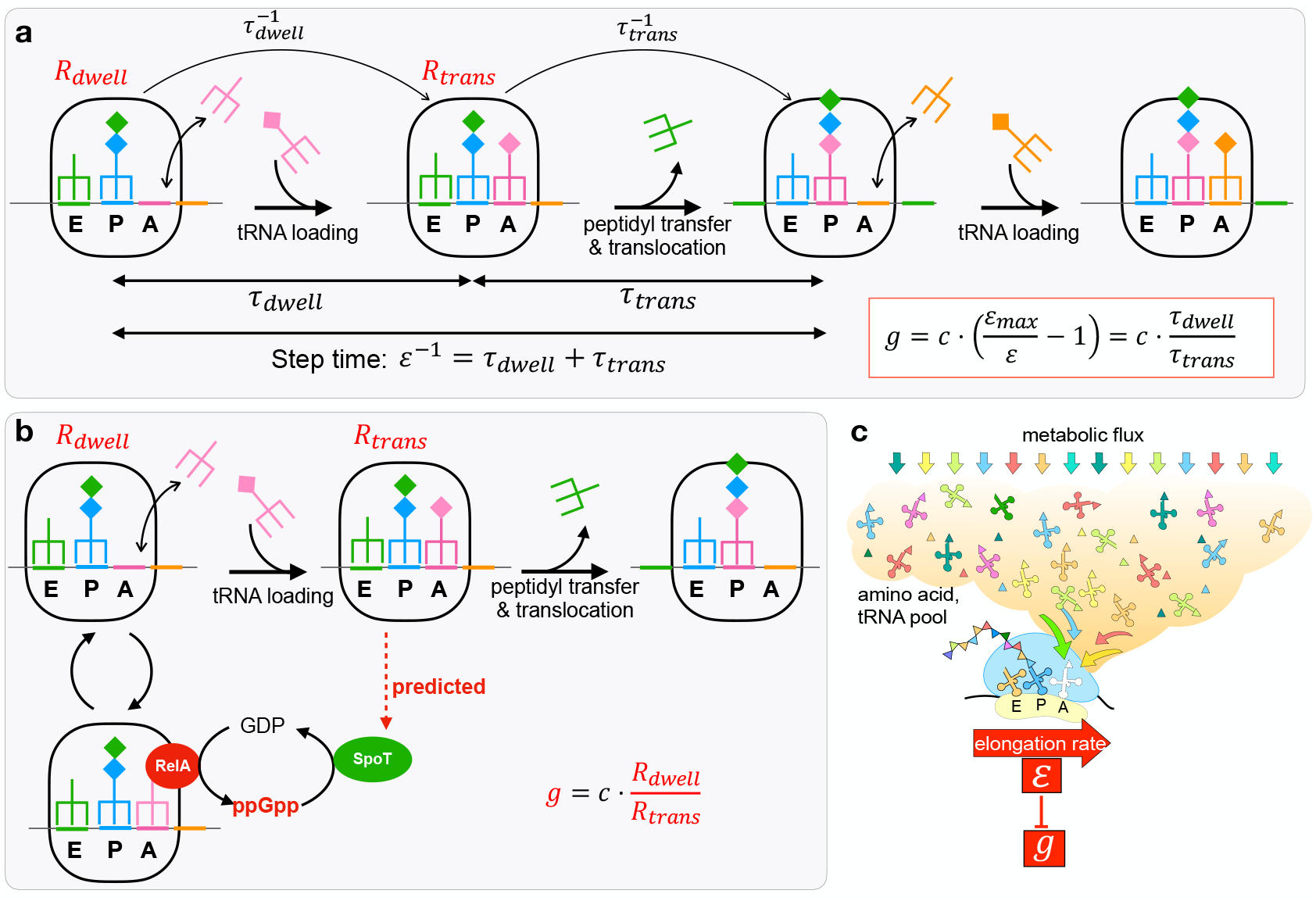
Sensing of the translational elongation rate by ppGpp. **a**, A cycle of translation elongation constitutes the loading of a cognate charged tRNA to the A site (taking time *τ*_*dwell*_), followed by peptidyl transfer and the translocation of mRNA/tRNA (taking time *τ*_*trans*_). The total time for one cycle, given by the reciprocal of the translational elongation rate *ε*, is thus given by *ε*^−1^ = *τ*_*dwell*_ + *τ*_*trans*_. *τ*_*trans*_ depends on the molecular properties of the translation machinery and *τ*_*dwell*_ depends on the concentration of uncharged tRNAs. Hence, long dwell times would lead to slow elongation speeds after a nutrient downshift owing to the increased uncharged tRNA levels, while the fastest elongation speed is obtained when *τ*_*dwell*_ → 0, in which case 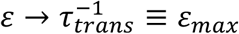. The empirical relation observed between ppGpp and the elongation rate *ε*, Eq. (1), can thus be alternatively written as *g* = *cτ*_*dwell*_/*τ*_*trans*_. According to the analysis of Supp. Note 2, the ratio of dwelling and translocation time is given by the ratio of the dwelling to translocating ribosomes, whose concentrations are *R*_*dwell*_ and *R*_*trans*_ as indicated in the figure. As the transition from the dwelling state to the translocation state occurs with rate 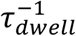 while the transition from the translocation to the dwelling state occurs with the rate 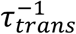, the condition of detail balance imposes that *τ*_*dwell*_/*τ*_*trans*_ = *R*_*dwell*_/*R*_*trans*_ regardless of the values of hundreds of molecular parameters underlying the translation process (Supplementary Note 2). It then follows that the empirical relation (1) can be obtained if, e.g., the synthesis of ppGpp is proportional to *R*_*dwell*_ and the hydrolysis of ppGpp is proportional to *R*_*trans*_. **b**, According to biochemical and structural studies^24,54,55,57,80^, ppGpp synthesis is activated when the A-site of the ribosome is loaded by a RelA-bound uncharged tRNA. This provides a mechanistic model for the control of ppGpp synthesis rate by *R*_*dwell*_, the concentration of dwelling ribosomes. Although this RelA-mediated synthesis activity would provide elevated levels of ppGpp in poor nutrient conditions, it is insufficient to generate the empirical relation described in Eq. (1); see Supplementary Figure 10 with details in Supplementary Note 2. Instead, the involvement of state-dependent ppGpp hydrolysis is predicted for a full account of the empirical relation. **c**, As charged tRNAs must be delivered to the ribosomal A-site to complete each step of translation, the distribution of dwelling and translocating ribosomes is dependent on the metabolic fluxes directed towards all the biosynthetic precursors needed for protein synthesis, represented here by the orange cloud: triangles, amino acids; clover leaves, tRNA; colored arrows, fluxes. The translational elongation rate is therefore a single quantity that reflects the combined status of the hundreds of diverse metabolic reactions underlying protein synthesis and cell growth (Supplementary Note 2). In this sense, the mechanism of ER-sensing described in panel b is a dimensional reduction technique employed by *E. coli* to convey the nutrient status by a single quantity, the level of ppGpp. The latter is further connected to the growth rate via the regulatory circuit of Box 1.

## Notes

### Competing Interest Statement

The authors have declared no competing interest.

### Summary of Updates

The revised version includes a detailed mathematical analysis which connects the observed relation between ppGpp and translational elongation rate to mechanisms of ppGpp synthesis and hydrolysis. Also added is an extended discussion on cellular strategy to implement the linear growth law between ribosome content and growth rate while maintaining a sizable pool of inactive ribosomes as reserve. This strategy challenges the common interpretation of the linear growth law as a manifestation of optimal resource allocation.

