## Supplementary Information for "Cellular perception of growth rate and the mechanistic origin of bacterial growth law"

### Supplementary Notes

##### Supplementary Note 1: Dependence of the steady-state model on $\varepsilon_0$ and $R_0$

In the main text, Eq. (5) described the relationship[a] between growth rate $\lambda$ and ppGpp level (or elongation rate via Eq. (1)) using parameter $a$ and $b$. Here we show that $a$ and $b$ can be expressed by $\varepsilon_0$ and $R_0$, defined as the elongation rate (ER) and ribosomal abundance, respectively, when the growth rate approaches zero. Also, we show that when $\varepsilon_0 : \varepsilon_{max} = 1:2$, the model yields simple dependences of $R$ and $g$ on $\lambda$.

In order for ER to be nonzero (with value $\varepsilon_0$) when the growth rate approaches zero, we must have $R = H$ (with value $R_0$) according to Eq. (5). Using Eqs. (1) and (3), we can express $a$ in terms of $\varepsilon_0$ and $R_0$:

$$a = c \cdot R_0 \cdot (h - 1), \tag{S1.1}$$

where we introduced $h \equiv \varepsilon_{max}/\varepsilon_0$ for convenience. Similarly, from Eqs. (1) and (4), the parameter $b$ can be expressed as

$$b = \frac{R_0}{c \cdot (h - 1)}. \tag{S1.2}$$

Conversely, we can express the two physiological parameters $R_0$ and $h$ in terms of the model parameters $a$ and $b$, with $R_0 = \sqrt{a \cdot b}$ and $h = 1 + c\sqrt{a/b}$.

Substitute Eqs. (3), (S1.1) and (S1.2) into Eq. (5), we obtain a relationship between $\lambda$ and $R$ with $h$ and $R_0$ being the parameters:

$$\lambda = \varepsilon_{max} \frac{R^2 - R_0^2}{R + R_0 \cdot (h - 1)}. \tag{S1.3}$$

Notice that when $h = 2$ (or $\varepsilon_0 : \varepsilon_{max} = 1:2$), Eq. (S1.3) is reduced to $\lambda = \varepsilon_{max} \cdot (R - R_0)$, or

$$R = R_0 + \lambda/\varepsilon_{max}, \tag{S1.4}$$

i.e., an exact linear relationship between $R$ and $\lambda$ with $1/\varepsilon_{max}$ being the slope of $R$-$\lambda$ plot.

[a] In writing down the relationship in Eq. (2) and Eq. (5), we assumed proteins are stable. At very slow growth, the effect of protein degradation should in principle be included. However, based on the measured protein degradation rates (2-3%/h in both steady-state growth and stationary phase)[15,16], this effect is negligible even at the smallest growth rate we studied here.

We can similarly work out the relation between $g$ and $\lambda$. From Eq. (S1.1) above, we can rewrite Eq. (3) of the main text as $R = R_0 c \cdot (h-1)/g$. Defining the value of $g$ as the growth rate $\lambda \to 0$ as $g_0$, we have $c \cdot (h-1) = g_0$, or

$$R = R_0 \cdot g_0/g. \tag{S1.5}$$

Substituting Eq. (S1.5) into Eq. (S1.3), we obtain

$$\lambda = \varepsilon_{max} R_0 \frac{(g_0/g)^2 - 1}{g_0/g + (h-1)}. \tag{S1.6}$$

For $h = 2$ (or $\varepsilon_0 : \varepsilon_{max} = 1:2$), Eq. (S1.6) is reduced to $\lambda = \varepsilon_{max} R_0 \cdot (g_0/g - 1)$, or

$$g = \frac{g_0}{1 + \lambda/(\varepsilon_{max} R_0)}, \tag{S1.7}$$

i.e., a simple hyperbolic dependence of the ppGpp level on the growth rate $\lambda$. Alternatively, $g^{-1}$ has a simple linear dependence on $\lambda$.

Lastly, we examine the relation between $R$ and $\varepsilon$. Inverting Eq. (1) of the main text, we have

$$\varepsilon = \frac{\varepsilon_{max}}{1 + g/c}. \tag{S1.8}$$

Further using Eq. (S1.5) and $g_0 = c \cdot (h-1)$, we obtain

$$\varepsilon = \frac{\varepsilon_{max}}{1 + (h-1) R_0/R}, \tag{S1.9}$$

which is generally of the Michaelis form, $\varepsilon_{max}$ being the maximal ER. For $h = 2$, the "Michaelis constant" becomes simply $R_0$.

**Supplementary Note 2: Molecular-level models linking Elongation Rate and ppGpp**

The observed relation between ppGpp levels and the elongation rate (Eq. 1 of main text) provides a constraint on the molecular mechanisms underlying ppGpp synthesis and degradation. Given the mechanistic view of translation kinetics detailed in Box 2 of the main text, the observed relation in Eq. 1 can be expressed as $g = c\tau_{dwell}/\tau_{trans}$, where $\tau_{trans}$ is the translocation time of charged ribosomes and $\tau_{dwell}$ is the dwell time of ribosomes waiting for the cognate charged tRNA. However, it remains to be established how the cell can sense these two timescales biochemically, and how the timescales in turn reflect the various tRNA charging levels that ultimately determine the elongation rate. In this note, we show that the cell can feasibly sense the ratio $\tau_{dwell}:\tau_{trans}$ through the ratio of ribosomal states, $R_{dwell}:R_{trans}$ where $R_{dwell}$ is the concentration of ribosomes in the dwelling state and $R_{trans}$ is the concentration in the translocation state. We show that the relation $\tau_{dwell}:\tau_{trans} = R_{dwell}:R_{trans}$ depends only on an assumption of flux balance for each tRNA species during the translation process. We then show how the ratio of ribosomal states is proportional to a weighted average of tRNA charging ratios, such that the cell can sense the limitation of any charged tRNA species via sensing the ratio $R_{dwell}:R_{trans}$. Finally, we show how the known mechanism of ppGpp synthesis via RelA cannot alone recapitulate the observed relation between ppGpp and the elongation. We propose that regulation of SpoT, either directly or indirectly, by translocating ribosomes is needed to reproduce the observed relation. The proposed regulation of SpoT also provides a feasible mechanism for the cell to sense $R_{dwell}:R_{trans}$.

<u>*Elongation timescales and translocating/dwelling ribosomes*</u>

In order to produce the empirically observed proportionality between $\frac{\varepsilon_{max}}{\varepsilon} - 1$ and the ppGpp level, $g$, the cell must be able to sense the ratio of the time scales $\tau_{dwell}$ and $\tau_{trans}$ detailed in Box 2. In correspondence with these two timescales, we assume actively translating ribosomes can be in one of two states: dwelling, with concentration $R_{dwell}$, and translocating, with concentration $R_{trans}$ such that $R_{act} = R_{dwell} + R_{trans}$. From global flux balance, it follows that the growth rate, $\lambda$, must balance both 1) average elongation rate, $\varepsilon$, multiplied by the active ribosomes, $R_{act}$, as well as 2) the translocation rate, $\varepsilon_{max}$, multiplied by the translocating ribosomes, $R_{trans}$, such that

$$\lambda = \varepsilon \cdot R_{act} = \varepsilon_{max} R_{trans}. \qquad \text{(S2.1)}$$

Rearranging Eq. (S2.1) we see that the ratio of the elongation rate to the maximum rate is equal to the fraction of active ribosomes that are translocating

$$\frac{\varepsilon}{\varepsilon_{max}} = \frac{R_{trans}}{R_{act}}. \qquad \text{(S2.2)}$$

As discussed in Box 2, the elongation rate decomposes into two timescales such that, $\varepsilon^{-1} = \tau_{trans} + \tau_{dwell}$, and the maximum elongation rate can be interpreted as and $\varepsilon_{max}^{-1} = \tau_{trans}$. By substituting these expressions, along with $R_{act} = R_{dwell} + R_{trans}$, into Eq. (S2.2), we have

$$\frac{\tau_{trans}}{\tau_{trans}+\tau_{dwell}}=\frac{R_{trans}}{R_{trans}+R_{dwelll}}, \tag{S2.3}$$

Simplifying this expression then yields Eq. (6) in the main text: $R_{dwell}\cdot\tau_{dwell}^{-1}=R_{trans}\cdot\tau_{trans}^{-1}$. As mentioned, Eq. (6) must hold to ensure detailed balance of fluxes between the two ribosomal states.

Eq. (S2.2) can also be re-arranged to solve for elongation rate in terms of $R_{trans}$ and $R_{dwell}$, such that

$$\varepsilon=\frac{1}{\tau_{trans}}\frac{R_{trans}}{R_{dwell}+R_{trans}}. \tag{S2.4}$$

Using Eq. (S2.4) we can then see that the empirical quantity of interest, $\frac{\varepsilon_{max}}{\varepsilon}-1$, can be expressed in terms of translocating and dwelling ribosomes as

$$\frac{\varepsilon_{max}}{\varepsilon}-1=\frac{\tau_{dwell}}{\tau_{trans}}=\frac{R_{dwell}}{R_{trans}}. \tag{S2.5}$$

Thus, $\frac{\varepsilon_{max}}{\varepsilon}-1$ -- which corresponds to the ratio of the dwell and translocation times -- can be obtained by the cell simply as the ratio of the dwelling ribosomes to the translocating ribosomes. Note that this relation is valid *instantaneously*. In particular, Eq. (S2.5) depends only on the ratio of ribosomes in different states, not on the overall abundance of the translating ribosomes.

*tRNA charging and elongation rate*

Past models[1,2] have derived the dependence of the translation elongation rate on the individual tRNA abundances and charging levels. Here we use these models to connect the elongation timescales, $\tau_{trans}$ and $\tau_{dwell}$, to the individual tRNA charging levels. In the past models, each active (i.e., translating) ribosome is assumed to be located at a specific codon associated with a given tRNA species, $i$. At a given moment, the active ribosomes are partitioned into *i* subspecies, with $N_{R,i}^{act}$ representing the number of ribosomes whose A-site resides at a codon associated with the $i^{\text{th}}$ tRNA species. The total number of active ribosomes is $N_R^{act}=\sum_i N_{R,i}^{act}$. The corresponding concentrations of active ribosomes are $R_{act,i}\equiv N_{R,i}^{act}/M_P$ and $R_{act}\equiv N_R^{act}/M_P$. The concentration of each tRNA species $i$ is denoted as $t_{tot,i}$ and can be further partitioned as either charged, of concentration $ta_i$, or uncharged, of concentration $t_i$, such that $t_{tot,i}=t_i+ta_i$. Active ribosomes are assumed to undergo reversible binding with both cognate charged and uncharged tRNA species, having codon-specific dissociation constants $\kappa_i^{ta}$ and $\kappa_i^{t}$ respectively. Binding to non-cognate tRNAs is neglected. Under the reversible binding model, active ribosomes associated with the $i^{\text{th}}$ tRNA can be further partitioned into three possible states based on the occupancy of their A-site: 1) bound with charged tRNA, of concentration $R_{ta,i}$ and referred to as "charged ribosomes", 2) bound with uncharged tRNA, of concentration $R_{t,i}$ and referred to as "uncharged ribosomes", or 3) not bound to either, of concentration $R_{o,i}$ and referred to as "open ribosomes". It then holds that $R_{act,i}=R_{ta,i}+R_{t,i}+R_{o,i}$. Using the reversible binding model, the fractional abundance of the charged, uncharged and open ribosome species can be written as

$$\rho_i^{ta} \equiv \frac{R_{ta,i}}{R_{act,i}} = \left(\frac{ta_i}{\kappa_i^{ta}}\right) \Big/ \left(1 + \frac{ta_i}{\kappa_i^{ta}} + \frac{t_i}{\kappa_i^t}\right), \tag{S2.6}$$

$$\rho_i^{t} \equiv \frac{R_{t,i}}{R_{act,i}} = \left(\frac{t_i}{\kappa_i^{t}}\right) \Big/ \left(1 + \frac{ta_i}{\kappa_i^{ta}} + \frac{t_i}{\kappa_i^t}\right), \tag{S2.7}$$

$$\rho_i^{o} \equiv \frac{R_{o,i}}{R_{act,i}} = 1 \Big/ \left(1 + \frac{ta_i}{\kappa_i^{ta}} + \frac{t_i}{\kappa_i^t}\right). \tag{S2.8}$$

To relate these ribosome fractions to the translational rate of the ribosomes, we start with the incorporation flux of the amino acid associated with the $i^{\text{th}}$ tRNA, denoted as $J_i$. The flux $J_i$ is proportional to the number of ribosomes associated with the $i^{th}$ charged tRNA such that $J_i = \varepsilon_{max}\rho_i^{ta} N_{R,i}^{act}$. The proportionality constant is the maximum specific elongation rate, $\varepsilon_{max}$, which is just the rate for ribosome translocation given that it is loaded with the cognate charged tRNA. Due to flux balance of individual amino acids in protein synthesis, the consumption of amino acids from each tRNA species must be balanced by the overall protein synthesis flux, $J_R$, i.e., $J_i = f_i \cdot J_R$ where $f_i$ is the fraction of all actively translated codons corresponding to the $i^{\text{th}}$ tRNA species. It will be convenient to express the fluxes in intensive quantities. The protein synthesis flux is generally defined as $J_R \equiv \frac{d}{dt} M_P = \lambda(t) \cdot M_P$ where $\lambda(t) = \frac{d}{dt} \ln M_P$ is the instantaneous growth rate. This leads to

$$f_i \lambda = \varepsilon_{max} R_{act,i} \rho_i^{ta}, \tag{S2.9}$$

which is valid not only in steady state but also during transient where $\lambda$ and $\rho$ may be strongly time-dependent.

To derive an expression for the elongation rate, we solve for $R_{act,i}$ from Eq. (S2.9)

$$R_{act,i} = \frac{\lambda}{\varepsilon_{max}} \frac{f_i}{\rho_i^{ta}}, \tag{S2.10}$$

and sum it over all tRNA species to obtain

$$R_{act} = \sum_i R_{act,i} = \frac{\lambda}{\varepsilon_{max}} \sum_i f_i / \rho_i^{ta}\ . \tag{S2.11}$$

The translation elongation rate is then obtained from its definition as

$$\varepsilon \equiv \frac{\lambda}{R_{act}} = \varepsilon_{max} \Big/ \sum_i \frac{f_i}{\rho_i^{ta}} = \varepsilon_{max} \Big/ \left[1 + \sum_i f_i \frac{\kappa_i^{ta}}{ta_i} \left(1 + \frac{t_i}{\kappa_i^t}\right)\right], \tag{S2.12}$$

where the 2nd equality follows from the definition of $\rho_i^{ta}$ in Eq. (S2.6), so that the elongation rate can be expressed in terms of the charged and uncharged tRNA species. Importantly, Eq. (S2.12) can be rearranged such that

$$\frac{\varepsilon_{max}}{\varepsilon} - 1 = \sum_i f_i \frac{\kappa_i^{ta}}{ta_i}\left(1 + \frac{t_i}{\kappa_i^t}\right), \tag{S2.13}$$

where the left-hand side is the quantity found empirically to be proportional to the ppGpp level (Eq. 1 of the main text) both in steady state and during transient shifts (Fig. 2d).

It is also useful to express Eq. (S2.13) in terms of the ribosomes in their different states, to make contact with the results derived above based on global flux balance. From Eq. (S2.6) and (S2.10), we have $R_{ta,i} = R_{act,i} \cdot \rho_i^{ta} = f_i \lambda / \varepsilon_{max}$, and $R_{ta} \equiv \sum_i R_{ta,i} = \lambda / \varepsilon_{max}$ such that $R_{ta,i} = f_i \cdot R_{ta}$. Further, from Eqs. (S2.6)-(S2.8), we have

$$R_{t,i} = R_{ta,i} \frac{\rho_i^t}{\rho_i^{ta}} = f_i R_{ta} \cdot \left(\frac{\kappa_i^{ta}}{ta_i} \cdot \frac{t_i}{\kappa_i^t}\right), \tag{S2.14}$$

$$R_{o,i} = R_{ta,i} \frac{\rho_i^o}{\rho_i^{ta}} = f_i R_{ta} \cdot \left(\frac{\kappa_i^{ta}}{ta_i}\right). \tag{S2.15}$$

Summing up each of the above expression over $i$ and using the shorthand $R_t \equiv \sum_i R_{t,i}$, $R_o \equiv \sum_i R_{o,i}$, we obtain

$$R_t + R_o = R_{ta} \cdot \sum_i f_i \cdot \frac{\kappa_i^{ta}}{ta_i}\left(1 + \frac{t_i}{\kappa_i^t}\right). \tag{S2.16}$$

Comparing the above relation with Eq. (S2.13), we obtain

$$\frac{\varepsilon_{max}}{\varepsilon} - 1 = \frac{R_t + R_o}{R_{ta}}, \tag{S2.17}$$

i.e., our central quantity of interest, $\frac{\varepsilon_{max}}{\varepsilon} - 1$, is given by the ratio of the total concentrations of ribosomes not bound to the charged tRNAs ($R_t + R_o$) and those bound to charged tRNAs ($R_{ta}$). We can thus identify the former group as "dwelling ribosomes" of concentration $R_{dwell}$ referred to in the main text and earlier in this note, and the latter group as "translocating ribosomes" of concentration $R_{trans}$, with Eq. (S2.17) being a mathematical derivation of Eq. (S2.5).

*Implications for ppGpp synthesis and degradation*

It is known experimentally that ppGpp synthesis by RelA requires the binding of RelA-bound uncharged tRNA to the ribosome[3]. Thus, the pool of ribosomes that RelA samples is $R_t$ in our classification. On the other hand, Eq. (S2.17) requires that the observed ppGpp pool to be proportional to $(R_t + R_o)/R_{ta}$. This implies that $R_o \ll R_t$, which occurs if $t_i \gg \kappa_i^t$, i.e., it is much favorably for the A-site to be occupied by tRNA, even if uncharged, compared to not occupied at all. In this limit, we have $R_{dwell} \equiv R_t + R_o \approx R_t$, and Eq. (S2.17) becomes

$$\frac{R_{dwell}}{R_{trans}} \approx \sum_i f_i \frac{\kappa_i^{ta}}{\kappa_i^{t}} \frac{t_i}{ta_i}\ , \tag{S2.18}$$

which is a key result of this Note. It shows that the ppGpp level, which is directly proportional to the ratio of dwelling to translocating ribosomes, senses a weighted average of the inverses of the tRNA charging ratios, $ta_i : t_i$, each of which is in turn dictated by the availability of the corresponding amino acid pool. Thus by sensing the ratio of the dwelling and translocating ribosomes, ppGpp is able to combine the charging levels of the many tRNA species and hence the availability of each amino acid into a single signal.

The ratio $R_{dwell}/R_{trans}$ also provides the cell with a convenient biochemical "handle" which can be used to regulate ppGpp synthesis and degradation. Considering a simple model for ppGpp dynamics,

$$\frac{dg}{dt} = \alpha - \beta g, \tag{S2.19}$$

which yields $g = \alpha/\beta$. Given Eq. (S2.18), our empirical observation $g \propto \frac{\varepsilon_{max}}{\varepsilon} - 1$ can be most conveniently explained if ppGpp is synthesized at a rate proportional to $R_{dwell}$, such that $\alpha \propto R_{dwell}$, and hydrolyzed at a rate proportional to $g \cdot R_{trans}$, such that $\beta \propto R_{trans}$. The hypothesized form of the synthesis rate is well-justified by the known mechanism of RelA activity as explained in Box 2 of the main text. The hypothesized form of the hydrolysis rate could arise if SpoT's hydrolysis activity is stimulated by ribosomes bound to charged tRNA, or during the translocation of ribosomes. Alternatively, SpoT hydrolysis activity could be auto-regulated by ppGpp levels, e.g., $\beta \propto R_{act}/(1+g)$.

Regardless of the form, our analysis suggests that the regulation of SpoT hydrolysis activity is practically a requirement in order to produce the empirical relation between elongation rate and ppGpp level. (An exception is if ppGpp synthesis is controlled by the ratio of $R_{dwell}$ and $R_{trans}$, which would be very difficult to implement molecularly and is not considered here.) If ppGpp synthesis via RelA was the only point of regulation, with $\alpha \propto R^{dwell}$ and $\beta$ being constant as proposed in Bosdriesz et al[2], then ppGpp levels would be proportional to $R_{dwell}$. Combining Eq. (6) from the main text, $R_{dwell} \cdot \tau_{dwell}^{-1} = R_{trans} \cdot \tau_{trans}^{-1}$, and the constraint $R^{act} = R^{trans} + R^{dwell}$, we can show that $R^{dwell} = R^{act} \cdot \tau_{dwell}/(\tau_{trans} + \tau_{dwell})$. It then follows that, if RelA is the only point of regulation,

$$g \propto R^{act} \frac{\tau_{dwell}}{\tau_{trans} + \tau_{dwell}}. \tag{S2.20}$$

In this form it is clear that RelA-exclusive regulation would pose a problem in slow growth conditions where $g$ is observed to be in high abundance, because at slow growth $R_{act} \to 0$ while $\tau_{dwell}/(\tau_{trans} + \tau_{dwell})$ is limited by saturation. This is shown in Supplementary Fig. 10a where we have plotted observations in steady state growth from the main text against predictions for unregulated SpoT hydrolysis computed from Eq. (S2.20). Without regulation of SpoT, ppGpp would be much lower than observed. Likewise, there would be problems during the transient condition shown in Fig. 1c, where ppGpp reaches ~8 times their initial level. In such transitions the number of translating ribosomes $R_{act}$ may not change significantly right after the shift. However, the factor $\tau_{dwell}/(\tau_{dwell} + \tau_{trans}) = 1 - \varepsilon/\varepsilon_{max}$ changes only ~3x, from $\varepsilon/\varepsilon_{max} \approx 0.75$ before the shift (Supplementary Fig. 1e) to $\varepsilon/\varepsilon_{max} \approx 0.25$ shortly after the shift. This is clearly shown in Supplementary Fig. 10b where we have plotted transition data from the main text against predictions from Eq. (S2.20), assuming $R_{act}$ does not change from pre-shift conditions. Without SpoT regulation, ppGpp is predicted to undershoot observed levels. These problems would only be exacerbated if RelA-mediated activation of ppGpp synthesis saturates according to a Michaelis-Menten relation (i.e. $g \propto R_t/(R_t + K)$), as is proposed in Bosdriesz et al[2]. In such a case, ppGpp levels could only be more limited in both steady state and transient conditions. Combined, the slow-growth and transient ppGpp levels suggest that regulation of RelA by uncharged tRNA at the ribosomal A-site is by itself insufficient, and some regulation by SpoT is needed. It is therefore not surprising to see the ppGpp level responding in a different way in the $\Delta relA$ strain. However, quantitatively understanding the dynamics of the latter would require also knowing how ppGpp level is set in the steady state in the $\Delta relA$ strain.

### Supplementary Tables

**Supplementary Table 1: Strains and growth conditions used in this study.**

| strain name | description | growth condition | growth rate ($h^{-1}$) |
|---|---|---|---|
| NCM3722[4,5] | Wild type *E. coli* K-12 | MOPS+ 20mM glucose | 0.96 |
| | | MOPS+20mM succinate | 0.68 |
| | | MOPS+40mM glycerol | 0.65 |
| | | MOPS+30mM acetate | 0.43 |
| | | MOPS+3mM mannose | 0.33 |
| | | MOPS+20mM aspartate + 10mM $NH_4Cl$ | 0.35 |
| | | MOPS 0.4% glycerol+10mM arginine | 0.32 |
| | | MOPS+20mM glutamate + 10mM $NH_4Cl$ | 0.13 |
| NQ1261[6] | Δ*ptsG* | MOPS+20mM glucose | 0.36 |
| | | MOPS+20mM glucose + 2μM Cm | 0.16 |
| | | MOPS+20mM glucose + 3μM Cm | 0.12 |
| HE838 (this work) | Δ*relA* | MOPS 20mM glucose | 0.99 |
| | | MOPS+4mM mannose | 0.42 |
| | | MOPS+20 mM aspartate+10mM $NH_4Cl$ | 0.34 |
| NQ1468[6] | Inducible LacZ-α with constitutively expressed LacZ-ω | MOPS+ 20mM glucose | 0.90 |
| NCM 3722 and HE838 under diauxic transition | | MOPS+2mM glucose+40mM glycerol | N.A. |

**Supplementary Table 2: Primers used in this study.**

| name | sequence | use |
|---|---|---|
| relA-P1 | cgatttcggcaggtctggtccctaaaggagaggacgatggttg cggtaagatgtgtaggctggagctgcttc | Chromosomal deletion of *relA* gene |
| relA-P2 | atatcaatctacattgtagatacgagcaaatttcggcctaactccc gtgcaattccggggatccgtcgacctg | Chromosomal deletion of *relA* gene |
| relA-ver-R | tacgctactgtggatcataaccctttc | Verification of deletion of *relA* gene |

#### Supplementary Figures

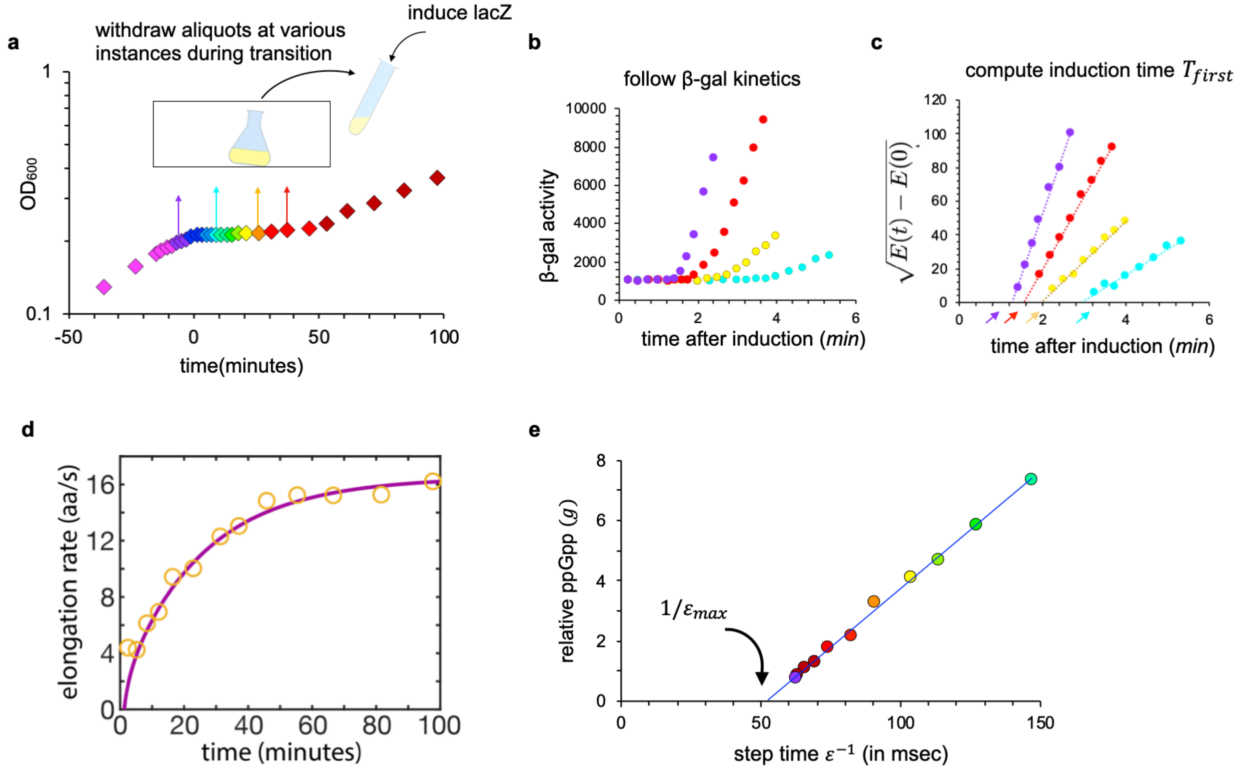

**Supplementary Figure 1. Elongation rate measurements during diauxic shift. a,** Scheme for measuring the instantaneous translation elongation rates $\varepsilon(t)$ during growth transition. At different time $(t)$ during the diauxic transition, 10 ml aliquots were removed into a fresh tube and the synthesis of the reporter protein LacZ was induced immediately by the addition of IPTG. **b,** LacZ induction curves for the four samples taken (indicated by the arrows in panel **a**). LacZ induction kinetics of each sample was followed by monitoring the β-galactosidase activity using 4-methylumbelliferyl-D-galactopyranoside (MUG) assay (see Methods and Materials for details). **c,** The square root of lacZ activity above basal level were plotted against induction time to obtain the lag time for the synthesis of the first LacZ molecule ($T_{first}$, shown by the arrows[7]). The translational elongation rates shown in Fig. 1 were obtained as $\varepsilon = L_{\text{LacZ}}/(T_{first} - T_{init})$ with $L_{\text{LacZ}}$ being the length of LacZ monomer (1024aa) and $T_{init}$ being the time taken for the initiation steps (10s across various nutrient conditions[6] and during transition; see **Supplementary Fig. 2**). **d,** Comparison of the elongation rate obtained from the naïve approach described above with a more detailed calculation using all of the LacZ induction data together. The yellow circle represents the naïve calculation of ER at the various time where samples were taken. The purple line represents $\varepsilon(t)$ calculated from solving the following integral equation

$$\int_{t_i+T_{init}}^{t_i+T_{first}(t_i)} \varepsilon(t)dt = \int_0^{t_i+T_{first}(t_i)} \varepsilon(t)dt - \int_0^{t_i+T_{init}} \varepsilon(t)dt = L_{\text{LacZ}}, \qquad \text{(E1)}$$

where $t_i$ is the time when IPTG is added. $T_{first}(t_i)$ denotes the lag time obtained from the LacZ induction curve when IPTG was added at time $t_i$. $T_{init}$ was taken to be 10s as explained in **Supplementary Fig. 2**. Taking the derivative of $t_i$ on both sides of Eq. (E1), we obtain

$$\varepsilon\left(t_i + T_{first}(t_i)\right) \times \left(1 + \frac{d}{dt_i} T_{first}(t_i)\right) = \varepsilon(t_i). \quad \text{(E2)}$$

By fitting $T_{first}(t_i)$ to an exponential function $y = a_1 \exp(a_2 x) + a_3$, we estimated $\frac{d}{dt_i} T_{first}(t_i)$ at each time point during the shift. For $t_i > 40$ min, since the change in $T_{first}$ is relatively small (circles in (d) represent the inverse of $T_{first}$), we Taylor-expanded $\varepsilon\left(t_i + T_{first}(t_i)\right)$ at $t_i$ and solved Eq. (E2) analytically with the boundary condition $\varepsilon(100 \text{ min}) = \varepsilon_{glycerol} = 16 aa/s$. For $t_i < 40$ min, we directly used Eq. (E2) to calculate $\varepsilon(t_i)$ from $\varepsilon\left(t_i + T_{first}(t_i)\right)$ at a later time numerically. Notice that the difference between the two ways of calculating the instantaneous ER is negligible. This is because the time scale of $T_{first}$, a few minutes as seen in panel (c), is overall much smaller than the time scale of the shift (~40 min). So we simply reported the result of the naive ER calculation through this work. **e,** Relative ppGpp level $g(t)$ obtained in Fig. 1c are plotted against the reciprocal of the elongation rate, or the step-time for ribosome advancement. $\varepsilon(t)$ values for the time points at which ppGpp levels were measured were estimated from fitting the naïve calculation of ER vs time in panel (d) to an exponential function. A linear fit of the data in panel (e) yields the x-intercept $1/\varepsilon_{max}$, with $\varepsilon_{max} = 19.4 \pm 1.4\ aa/s$.

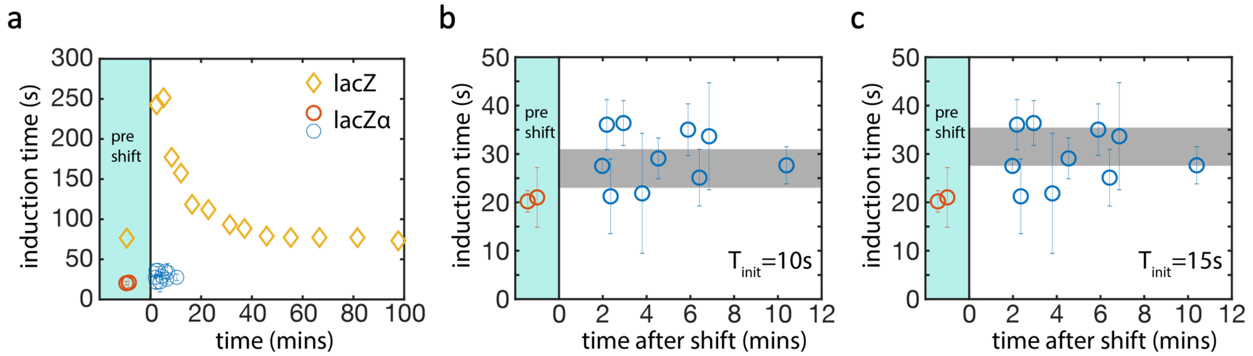

**Supplementary Figure 2. Determination of the translational initiation time.** The induction time of lacZ monomer ($T_{first}$ shown in **Supplementary Fig. 1c**) during the shift were plotted as yellow diamonds in panel (**a**). To obtain the contributions of translational initiation towards this induction time, we also determined the induction kinetics of a short LacZ alpha fragment ($LacZ\alpha$, the N-terminal 1-90aa of lacZ) before and right after the shift as described in Dai et al[6]. The induction time obtained, $T_{first,LacZ\alpha}$, were plotted as red and blue circles respectively. The error bars indicated the 95% confidence interval from fitting and technical repeats. The induction time obtained can in principle be used to estimate the translational initiation time, $T_{init} = T_{first,LacZ\alpha} - L_{LacZ\alpha}/\tilde{\varepsilon}$, where as a first estimate of the elongation rate, we used $\tilde{\varepsilon} = (L_{lacZ} - L_{lacZ\alpha})/(T_{first,LacZ} - T_{first,LacZ\alpha})$ assuming the same initiation time for LacZ and LacZ alpha, with $T_{first,LacZ\alpha}$ used for $T_{init}$ of LacZ$\alpha$. This was done for the pre-shift condition, with the result $T_{init} = 15s \pm 6s$ (error from uncertainty in $LacZ\alpha$ induction time), consistent with previous measurements[6]. To estimate the initiation time immediately following the shift, we looked more closely into the induction time data collected, shown as the blue circles in panels (**b**) and (**c**). Due to the uncertainties in $LacZ\alpha$ induction time determination, together with the uncertainty in determining the precise shift time (i.e., time "0") between the strains expressing LacZ and LacZ$\alpha$, it is difficult to calculate the initiation time based on the difference $T_{first,LacZ} - T_{first,LacZ\alpha}$ while $T_{first,LacZ}$ varied by strongly in the first 10 min. Instead, we assumed a constant initiation time ($T_{init}$) and predicted the range of $LacZ\alpha$ induction time given measured $LacZ$ induction time in this 10min-window (160s-250s). The grey bar in **b** and **c** shows the predicted range for $T_{first,LacZ\alpha}$ assuming $T_{init} = 10s$ and $T_{init} = 15s$, respectively. Comparing with our measurements, it suggest the initiation time in transition is within the range of 10s-15s, similar to the initiation time in pre-shift condition. For simplicity, we used $T_{init} = 10s$ for both pre- and post-shift throughout this work. This does not mean that the initiation time is not affected during the transition, but that our method is too coarse to resolve differences in the initiation time.

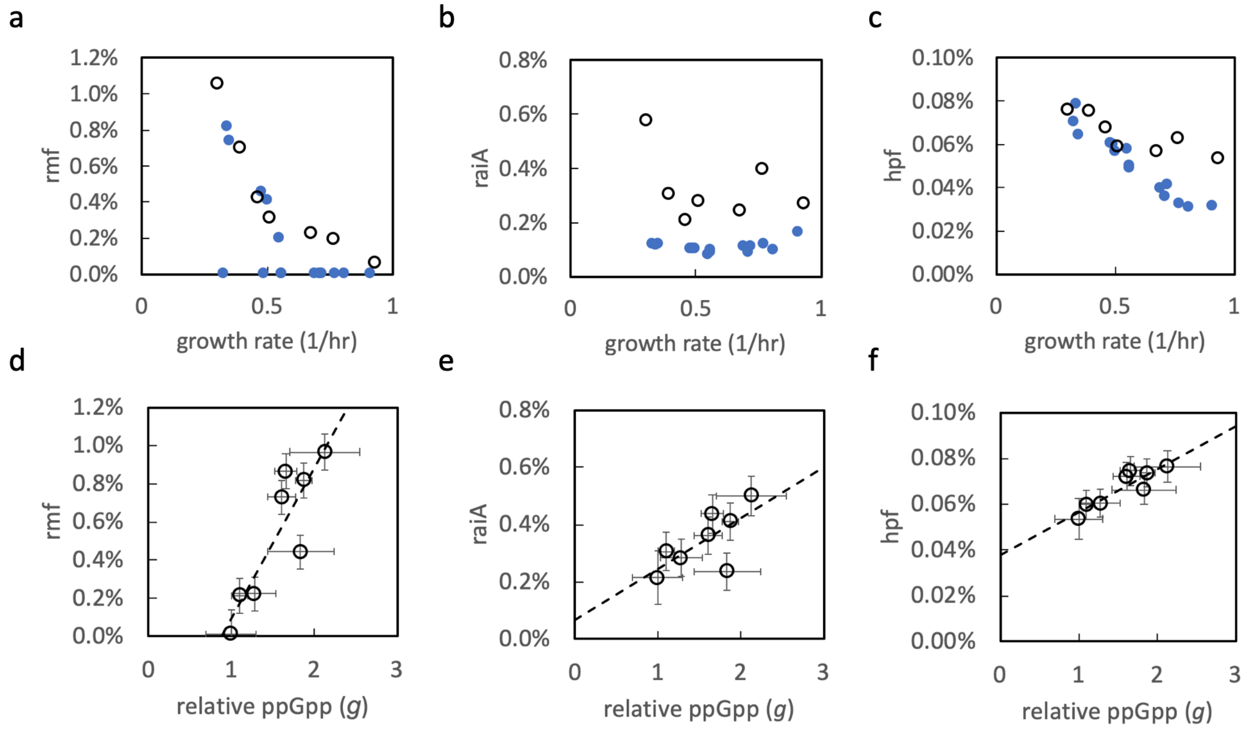

**Supplementary Figure 3: Relation between the ribosome-sequestering proteins and the ppGpp levels. a,** The number of mRNA (black circles) and protein (blue circles) as a percentage of the total number of mRNA and protein, respectively, for the ribosome remodeling factor Rmf obtained during steady state growth are plotted against the growth rate for a number of cultures under varying degrees of carbon limitation. Data for mRNA abundance is obtained from Balakrishnan et al[8] and for protein abundance is obtained from Mori et al[9]. **b,** Same as panel a, but for the mRNA and protein of the ribosome associated inhibitor gene *raiA* [10]. **c,** Same as panel a, but for the mRNA and protein of the gene encoding the hibernation promoting factor *hpf* [11]. For each case shown in panels a-c, the protein and mRNA abundances approximately match each other within 2-fold, as is typical of many genes expressed during exponential growth. For Rmf, the protein level showed zeero for a number of conditions due to lack of detected peptides. **d-f,** The fractional number abundances of the mRNA (black circles) of *rmf*, *raiA* and *hpf* are plotted against the relative ppGpp of the corresponding conditions. The black dash lines show the linear fit of the data. To directly compare the expression of *rmf, raiA* and *hpf* with ppGpp levels, we fitted the mRNA vs GR data in panel a-c with 3rd-degree polynomials and estimated their mRNA levels at the same growth rates for which ppGpp was measured. The horizonal error bars in panels d-f comes from ppGpp measurements, while the vertical error bars come from the polynomial fitting (95% prediction interval).

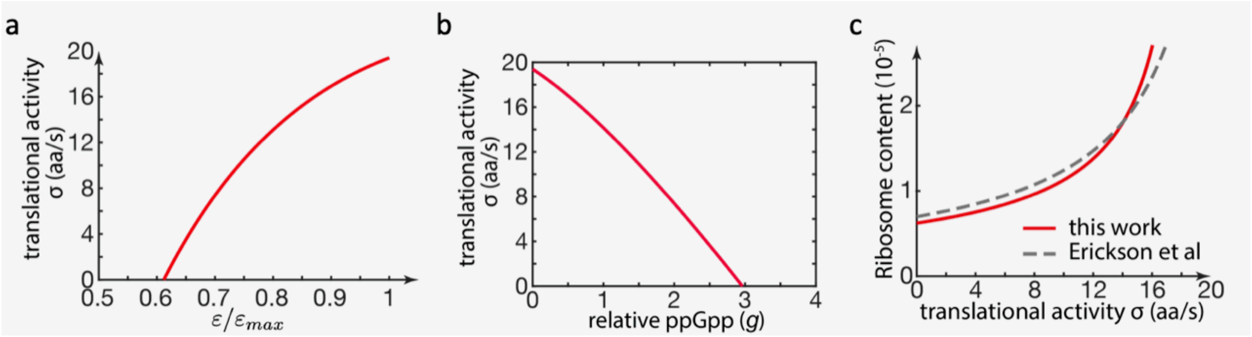

**Supplementary Figure 4. a,** Translational activity $\sigma$, defined as $\lambda/R$, and thus given by $\varepsilon \cdot (R - H)/R$, is plotted against normalized elongation rate for the best-fit model parameters used in Fig. 3. **b,** Translational activity $\sigma$ plotted against the relative ppGpp level $g$. The one-to-one relationship between the translational activity and the ppGpp level justifies the use of the translational activity as the dynamic variable in the kinetic model of diauxic transition developed by Erickson et al[12]. **c,** The plot of ribosomal content vs $\sigma$ used by Erickson et al[12] as the regulatory function for ribosome biogenesis is shown as the grey dashed line. The same relation according to the model in Fig. 3 is shown as the solid red line. The two lines are very similar, indicating that Erickson et al[12] correctly inferred the regulatory function even though the relation between $\sigma$ and $g$ (panel **c**) was not known at the time.

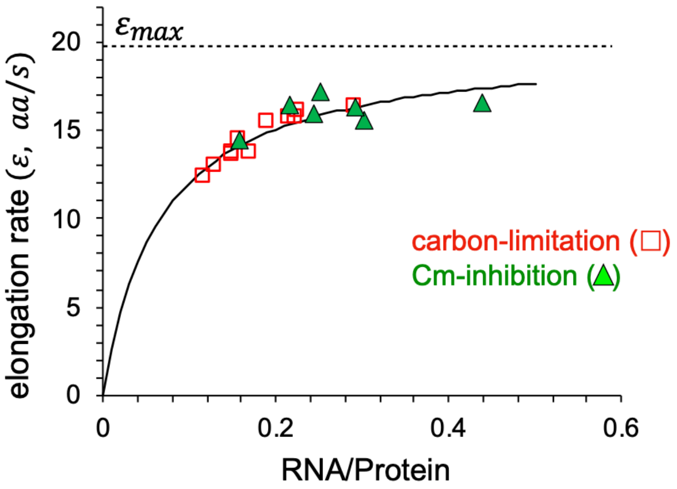

**Supplementary Figure 5: Estimating the maximum translation elongation rate in steady-state growth.** Translation elongation rates measured in the carbon-limited and chloramphenicol-treated cultures are plotted against the ratio of RNA and protein measured in the same conditions. The data are collectively fitted to the Michaelis-Menten relation $\varepsilon = \varepsilon_{max} \cdot r/(k_m + r)$ where $r$ is the RNA-protein ratio. The best-fit gives $\varepsilon_{max} = 20.0 \pm 1.9 aa/s$ and $k_m = 0.066$. The maximum elongation rate $\varepsilon_{max}$ estimated this way is indistinguishable from that estimated from the relation between ppGpp and the elongation rate during growth transition (see **Supplementary Fig. 1e**).

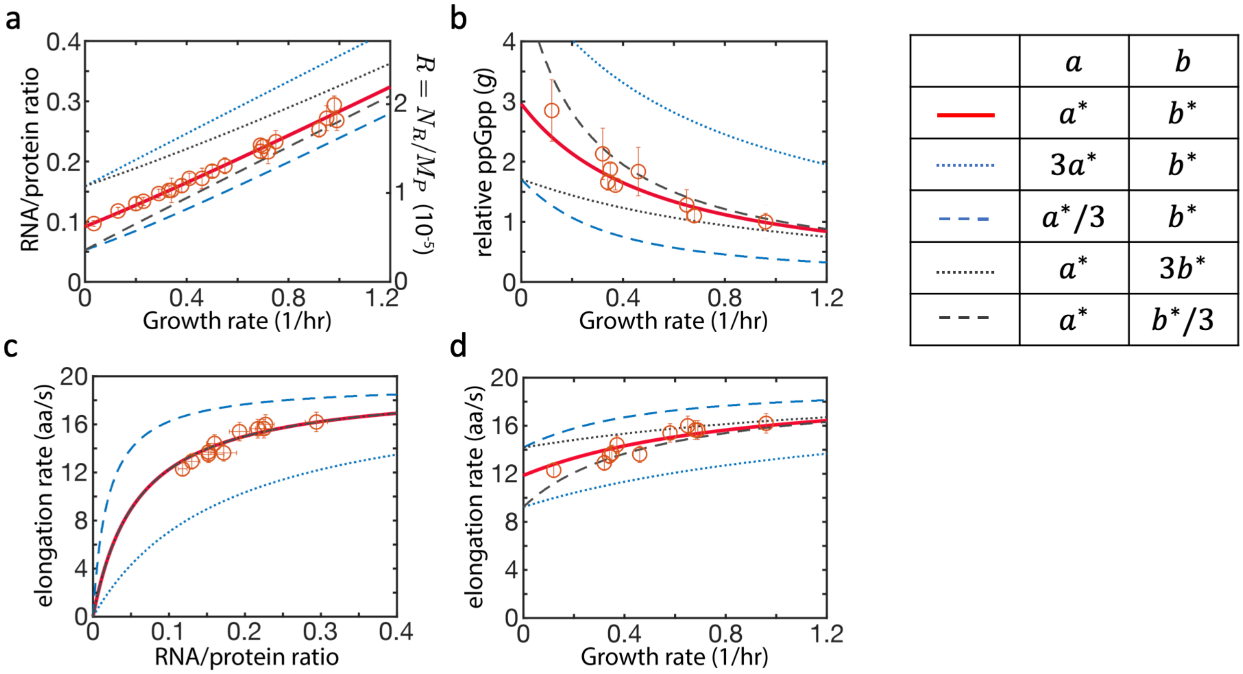

**Supplementary Figure 6: The effect of parameter choices on model output.** The measured data points (red open circles) and the best-fit model outputs (solid red lines) in panels a-d are the same as those shown in **Fig. 3b-e**. When keeping parameter $a$ the same as the best-fit value $a^*$ while varying $b$ to be 3-fold larger or smaller, the model gave predictions shown as the dotted and dash black lines, respectively. When keeping parameter $b$ the same as the best-fit value $b^*$ and varying $a$ to be 3-fold larger or smaller, the model gave predictions shown as the dotted and dash blue lines, respectively; see the legends table. The results show that model outputs are comparatively more sensitive to the choice of the parameter $a$ than $b$. Fortunately, the parameter $a$ involved in the relation between ppGpp and the ribosome content (see Eq. (3)) is well established by **Fig. 3a**.

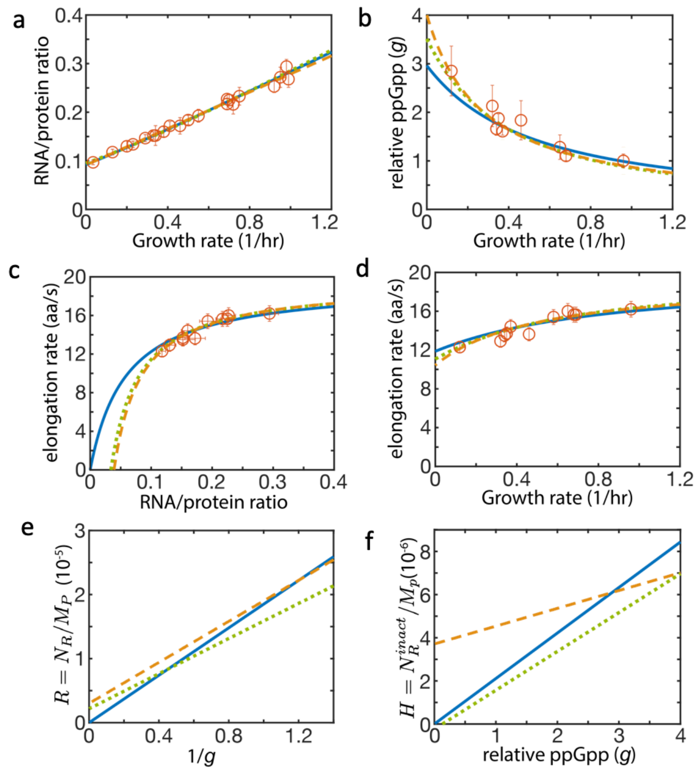

| | $g_H = 0$, $g_R = \infty$, $\frac{rRNA}{RNA}$ fitted | $g_H$ fitted, $g_R$ fitted, $\frac{rRNA}{RNA}$=0.86 | $g_H$ fitted, $g_R$ fitted, $\frac{rRNA}{RNA}$ fitted |
|---|---|---|---|
| $a$ | $1.85\times10^{-5}$ | $1.37\times10^{-5}$ | $1.6\times10^{-5}$ |
| $b$ | $2.11\times10^{-6}$ | $1.8\times10^{-6}$ | $8.25\times10^{-7}$ |
| $g_R$ | | 6.3 | 5.3 |
| $g_H$ | | −0.13 | 4.5 |
| $\frac{rRNA}{RNA}$ | 0.92 | | 1.03 |

$$R(g) = a \cdot (1/g + 1/g_R) \quad \text{---(E3)}$$

$$H(g) = b \cdot (g + g_H) \quad \text{---(E4)}$$

**Supplementary Figure 7. The effect of constant offsets in the regulatory functions on model output.** Here we investigate the effect of altered forms of regulatory functions from the simplest forms assumed in Eqs. (3) and (4). The altered forms are shown as Eqs. (E3) and (E4) below the legend table on the right, with constant offsets parameterized by $g_R$ and $g_H$. **a-d** show the same four outputs as in Supplementary Figure 6 with different modeling results. The table on the right compares the best-fit parameter values under different model settings. The solid blue line is the same as that shown in **Fig. 3** (where offsets in $R(g)$ or $H(g)$ are not considered, left column of the legend). The orange dash lines are the best-fit of the model with $g_R$ and $g_H$ treated as fitting parameters also. Results of the fits are listed in the right column of the legend table. While the fits may be marginally better, one aspect of the fitting parameter becomes quite unreasonable: the relation between the ribosome content ($N_R/M_P$) and the RNA-protein ratio is determined by the fraction of ribosome RNA within all RNA (the last row in legend table). In the case where all parameters are fitted (right column), the rRNA-RNA ratio is ~1, which is impossible given that tRNA comprises ~10% of the total RNA. In comparison, the simplest model not considering offsets gave an rRNA-RNA ratio of 92% (left column) which is closer to the accepted range. If we fix the rRNA-RNA ratio to 86%, estimated previously for a few growth conditions[13,14] , the best-fit model outputs are shown as the green dotted lines, with the best-fit parameters shown in the middle column. For the ease of assessing the effect of these offsets on the form of the regulatory functions $R(g)$ and $H(g)$, we display these forms for the 3 cases considered in panel **e** and **f**, using the same line styles as those indicated in the legend table. We see that little difference is made to the model outputs despite substantial changes to the forms of the regulatory functions, thus indicating that these results are robust to the exact forms of the regulatory functions assumed.

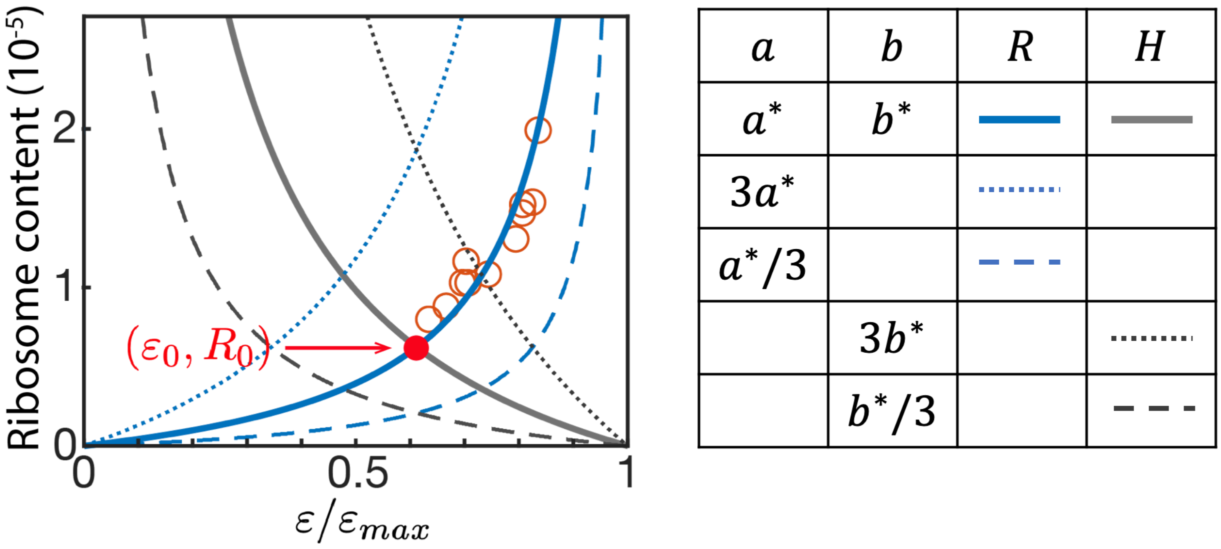

**Supplementary Figure 8.** Ribosome content are plotted against normalized elongation rate for various model parameters. Using the best-fit parameters in Fig. 3 (labeled as $a^*$ and $b^*$ in legend table), total ribosome content (solid blue line) increases with elongation rate while inactive ribosome content (solid grey line) decrease with elongation rate. Their intersection (marked by the filled red circle) represents the elongation rate and ribosome content while growth rate approaches zero, i.e., $\varepsilon_0$ and $R_0$, respectively. When the parameter $a$ is changed to 3-fold smaller or larger, the relation between total ribosome content and elongation rate is changed to the blue dashed or dotted lines, respectively. When the parameter $b$ is changed 3-fold smaller or larger, the relation between total ribosome content and elongation rate is changed to the black dashed and dotted lines, respectively. As shown in the figure, the different parameters result in different values of the intersection point and hence different values of $(\varepsilon_0, R_0)$.

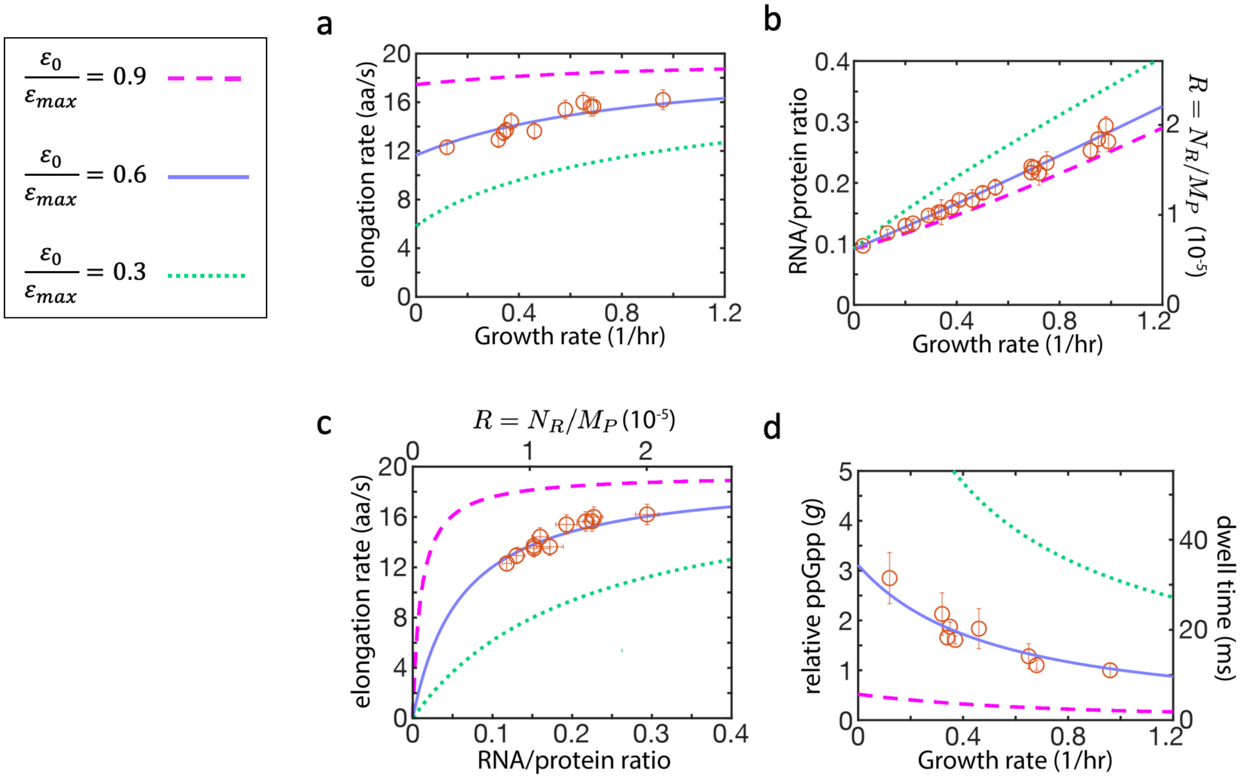

**Supplementary Figure 9. Dependence of model output on the ratio $\boldsymbol{\varepsilon_0 : \varepsilon_{max}}$.** As shown in Supplementary Note 2, the model defined by Eqs. (1) and (5) has a particularly simple solution for $\varepsilon_0 = \varepsilon_{max}/2$, where $\varepsilon_0$ is the elongation rate as the growth rate $\lambda \to 0$. The best-fit output of the model has $\varepsilon_0 : \varepsilon_{max} \approx 0.6$, which is close to the simple linear solution, thus rationalizing the approximate linear correlations observed in Fig. 3. In this figure, we show the model output for choices of the parameters $a$ and $b$ such that $\varepsilon_0 : \varepsilon_{max} = 0.3$ and 0.9 (dotted and dashed lines, respectively). The red open circles are the same data as those shown in Fig. 3. Deviations from the linear growth law are clearly seen for the expected RNA/protein ratio in panel b, with the high ratio of $\varepsilon_0 : \varepsilon_{max}$ (dashed purple lines) exhibiting reduced ribosome content. In principle, large ER and reduced RNA/protein ratio are advantageous. However, this would lead to very low dwell time for ribosome on A-site as seen in panel d (right vertical axis, computed according to $g = c\ \tau_{dwell}/\tau_{trans}$): 10-30 ms for the best-fit parameters, and < 5ms for the purple dashed line with ER maintained above 90% of $\varepsilon_{max}$.

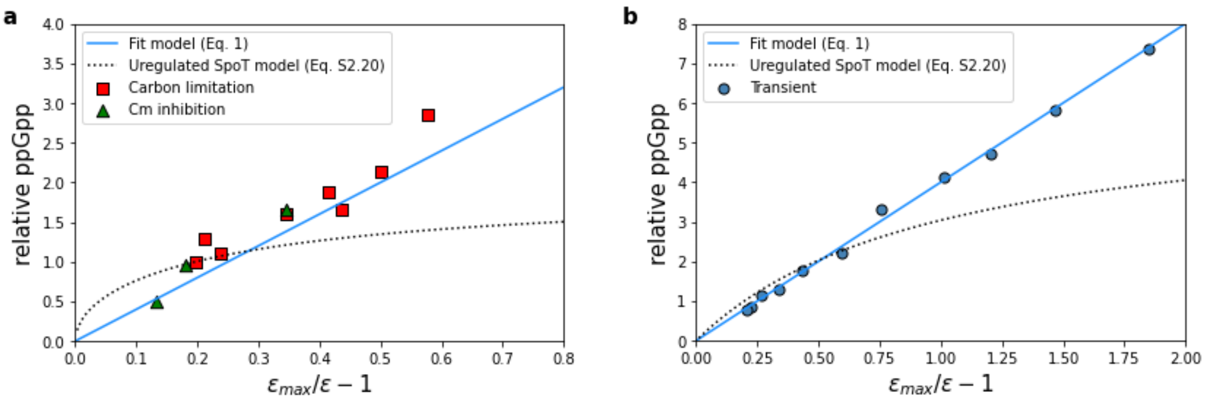

**Supplementary Figure 10. Comparing predictions for unregulated hydrolysis to the observed relation between elongation and ppGpp. a,** Observed ppGpp levels for steady state growth under carbon limitation and Cm inhibition (Fig. 2d) are compared to the predicted ppGpp levels for unregulated SpoT hydrolysis derived in Eq. (S2.20). Using Eq. (S2.20), $\tau_{dwel}/(\tau_{dwell} + \tau_{trans})$ is computed from the elongation rate as $1 - \varepsilon/\varepsilon_{max}$. The ppGpp level, $g$, is then solved assuming $R_{act}$ is the difference between $R(g)$, main text Eq. (3), and *H(g)*, main text Eq. (4). The proportionality constant for Eq. (S2.20) has been set so that the ppGpp level in the glucose reference condition is 1. The fit phenomenological model, main text Eq. (1), is shown for comparison with $\varepsilon_{max} \approx 19.4\ aa/s$ and $c \approx 4.0$. **b**, Observed ppGpp levels during the growth transition (Fig. 1e) are compared to predictions for unregulated SpoT hydrolysis computed from Eq. (S2.20). Here $R^{act}$ is assumed to remain at pre-shift levels, and the same proportionality constant is used from panel (a). The fit phenomenological model, main text Eq. (1), is again shown for comparison.

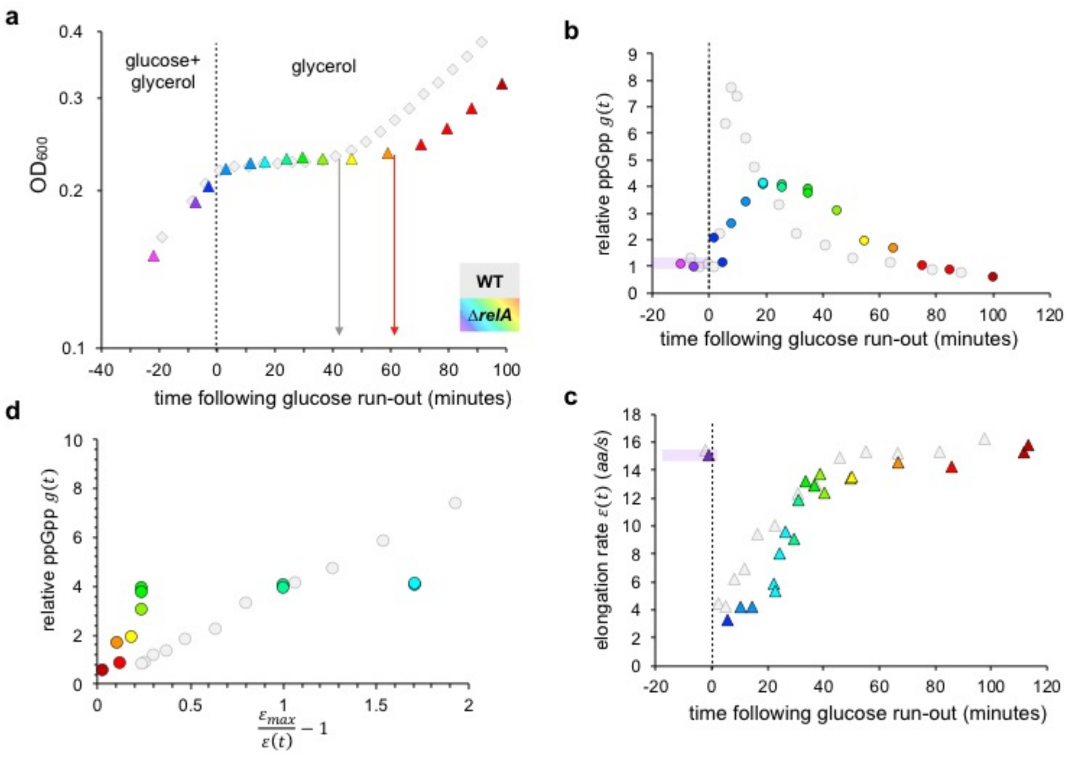

**Supplementary Figure 11. Relation between the ppGpp level and elongation rate for the Δ*relA* strain during diauxic shift. a,** Growth kinetics of the Δ*relA* strain (colored triangles) and the wild type (grey circles, same as in **Fig. 1a**) during the glucose-to-glycerol diauxic transition. Vertical arrows indicate the duration of the lag period following glucose depletion before growth resumes on glycerol. Color-scheme corresponding to individual samples for Δ*relA* is shared across panels in this figure. **b,** ppGpp levels measured for the Δ*relA* strain (colored) and the wild type (grey, same as those in **Fig. 1c**), relative to the WT ppGpp level in glucose minimal medium, during the glucose-to-glycerol transition. Note that pre-shift ppGpp levels of the two strains are indistinguishable. **c,** Instantaneous translation elongation rate measured for the Δ*relA* strain (colored) and the wild type (grey, same as those in **Fig. 1d**) during the glucose-to-glycerol transition. **d,** The relation between ppGpp level and the instantaneous elongation rate for the Δ*relA* strain (colored) and the wild type (grey, same as those in **Fig. 1e**) during the glucose-to-glycerol transition.

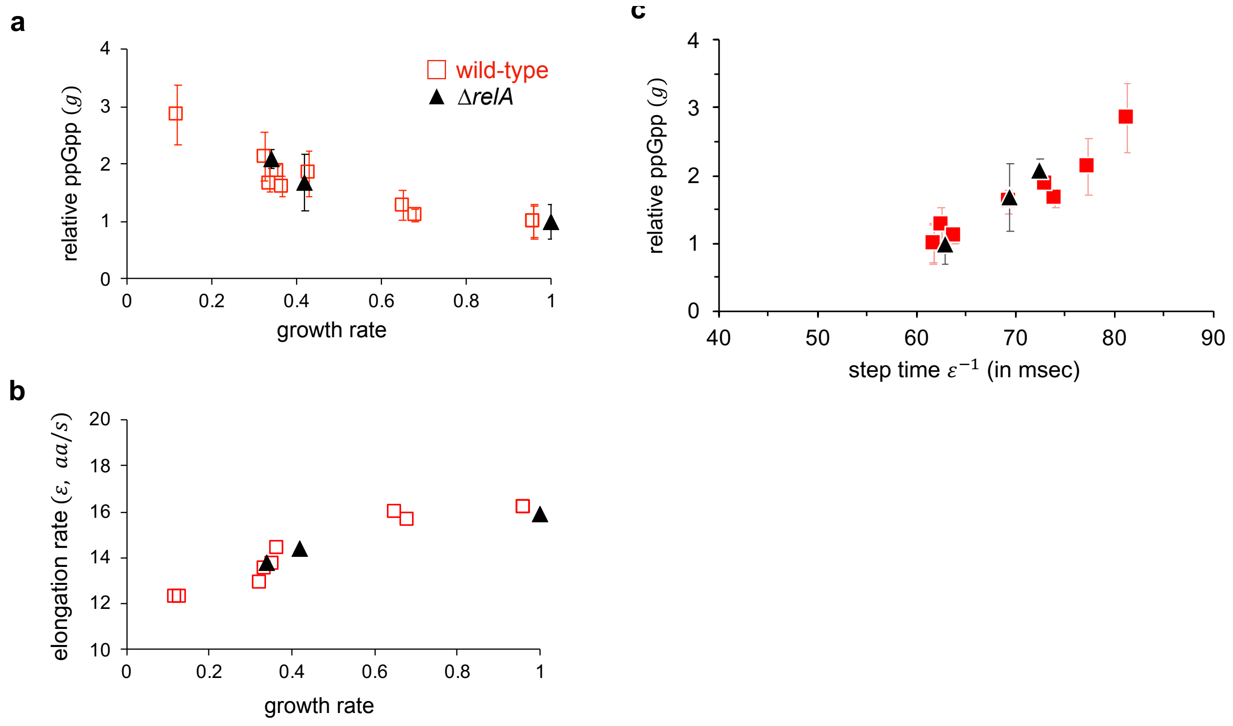

**Supplementary Figure 12. Relation between the ppGpp level and elongation rate for the Δ*relA* strain in steady-state growth. a,** ppGpp levels relative to that in the reference growth condition (wild type grown in minimal glucose medium) are plotted against the steady state growth rates for wild-type (red) and Δ*relA* (black) grown in various carbon sources (See **Supplementary Table 1**). Error bar represents the uncertainty in the linear fit over four measurements; see **Supplementary Fig. 13**. **b,** Translation elongation rates are plotted against the steady state growth rates for wild-type (red) and Δ*relA* (black) grown in various carbon sources (See **Supplementary Table 1**). **c,** Steady-state relative ppGpp levels are plotted against the reciprocal of the translational elongation rate for wild-type (red) and Δ*relA* (black) grown in the same conditions. Both the ppGpp levels and the elongation rates in steady state growth are not affected by *relA* deletion, unlike the case during diauxic transition shown in **Supplementary Fig. 11**.

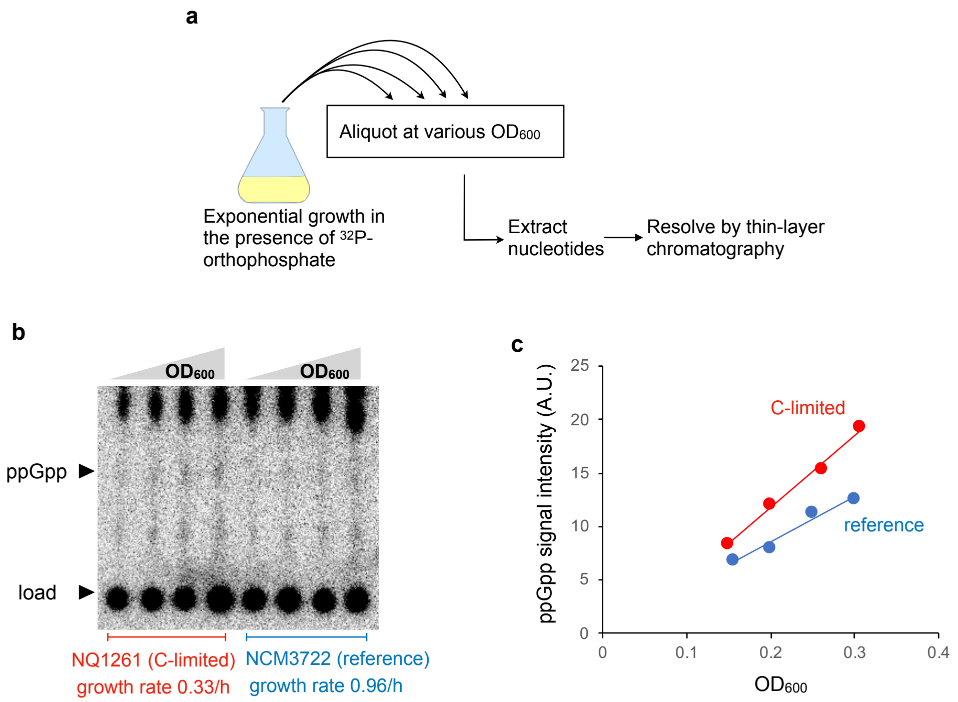

**Supplementary Figure 13: Measurement of ppGpp levels in steady-state growth conditions.** **a,** Scheme for ppGpp measurement under balanced growth. *E. coli* strains were grown in either glucose minimal media (reference condition, growth rate = 0.96/h) or various poor carbons at different steady-state growth rates. At various $OD_{600}$ values during the exponential growth phase of each culture, aliquots were withdrawn, nucleotides were extracted and ppGpp was resolved using TLC. **b,** An example of ppGpp measurement for the wild type strain (reference condition, blue) and NQ1261, a Δ*ptsG* strain defective in glucose uptake, both grown in glucose minimal medium (red). NQ1261 strain is used to create reduced growth on glucose (see **Methods**). Nucleotides extracted at four different $OD_{600}$ were spotted on the bottom of TLC plates and ppGpp was resolved. **c,** ppGpp from the wild type and NQ1261 strains are plotted against $OD_{600}$. The slope of each plot gives the ppGpp level (per $OD_{600}$) for that strain and condition. In **Fig. 2** of the main text, we report the ppGpp level relative to that in glucose steady state. In this case, the relative ppGpp level of NQ1261 strain grown in glucose is just the ratio of the slope of the red line to that of the blue line. Error bar in the estimate of ppGpp was taken to be the uncertainty in the slopes of the linear fit.
